# Mapping Four Decades of Microbial Waste Valorization Research in Africa through a Systematic Review and Bibliometric Analysis

**DOI:** 10.1101/2025.10.21.683771

**Authors:** Ndukwe K. Johnson, Dauda Wadzani Palnam, Peter Abraham, Israel Ogwuche Ogra, Okechukwu Kalu Iroha, Emohchonne Utos Jonathan, Elkanah Glen, Samson Usman, Seun Cecilia Joshua, Dogara Elisha Tumba, Mela Ilu Luka, Morumda Daji, Dasoem Naanswan Joseph, Grace Peter Wabba, Mercy Nathaniel, Friday Okibe, Zainab Kasim Mohammed, Umezuruike Linus Opara

## Abstract

The annual Africa waste production has been predicted to triple by 2050. Due to lack of proper waste disposal plans and services in many African countries, wastes are indiscriminately dumped leading to environmental and human health issues. Microorganisms with their complex biochemical functionaries play a significant role in waste valorization. Research in the application of microorganisms in waste conversion has been ongoing globally. However, there is no information on the extent of research on this subject with regards to Africa using bibliometric and systematic review approaches. This study was aimed at understanding the extent of research in microbially-driven waste valorization in Africa to identify research gaps and future perspectives. Bibliographic and systematic literature review (SLR) methodology guided by the Preferred Reporting Items for Systematic Reviews and Meta-Analyses (PRISMA) were used for this study. Bibliographic data (752) was extracted from Scopus database. Thereafter, 264 data were excluded and the remaining 488 data were analysed using Biblioshiny, Vosviewer and Excel analytical tools. Nigeria with 369 published articles ranked above other countries from 1984-2024. However, using the number of citations and single country publication (SCP) South Africa dominated with 3233 and 75% SCP. Themes like wastewater treatment, composting, recycling, and microorganisms evolved to biodegradation, biomass, microbial community, anaerobic digestion etc. The findings suggest that emerging themes are still revolving around biomass and anaerobic digestion, as more research is still on wastewater treatment and management, recycling and fermentations in waste valorization. This knowledge contributes to future research designing and gap identification in the area of waste valorization in Africa.

**Graphical abstract:** 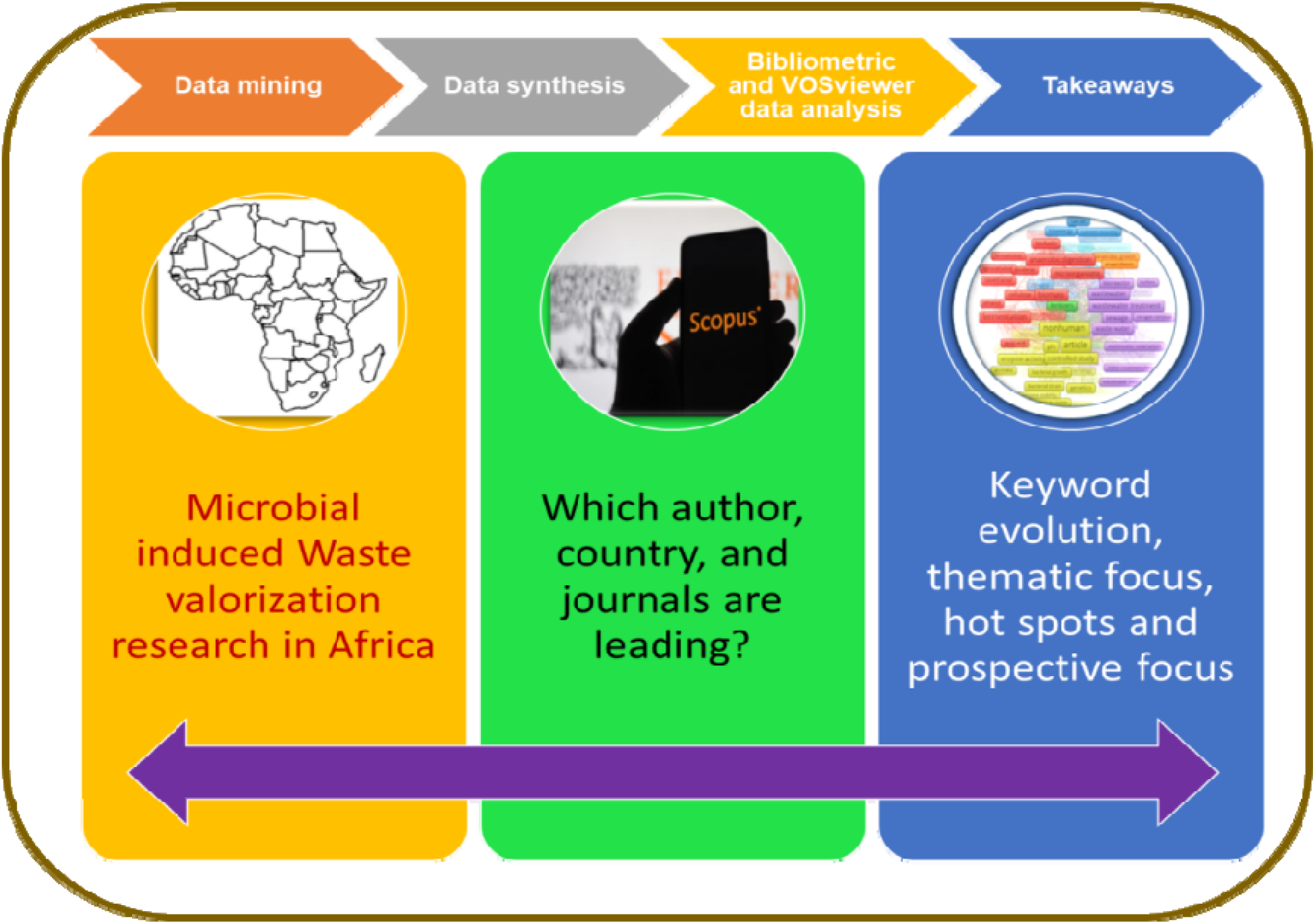

## 1.0 Introduction

Waste generation management is one of the biggest environmental management issues facing the globe today. Waste generation is on the rise yearly due to growth in the economy and unsustainable consumption and manufacturing practices (Dauda *et al*., 2023). The UNEP (2024) report projected that between 2020 and 2050, municipal solid waste (MSW) generation alone will increase by 56% within a generation or less, meaning that there will be a rise of MSW from 2.1 billion tonnes to 3.8 billion tonnes. The annual Africa waste production is expected to triple by 2050, from 174 million tonnes in 2016 to over 516 million tonnes (Tomita et al., 2020). According to available data, Africa alone produced 125 million tonnes of municipal solid waste (MSW) annually in 2012, with 81 million tonnes (65%) coming from sub-Saharan Africa (Scarlat et al., 2015); this is anticipated to increase to 244 million tonnes annually by 2025 (Godfrey et al., 2020). Kumari and Raghubanshi (2023) highlighted that waste generation management problems are more acute in developing and underdeveloped countries, as waste generation surpasses the population growth and economic development, making waste management a more critical issue. This wastes from both industries and homes are often dumped indiscriminately on the road, drainage systems, in bushes, around houses (both residential and offices), within streets, etc. In towns and villages due to lack of proper waste disposal plans and services (Dladla et al., 2016). Also, Africa houses up to 38% of the largest landfills in the world (UNEP, 2015); more than Asia, Latin America and Caribbean (Mwangomo, 2019). Diseases such as malaria, typhoid, cholera, dengue, yellow fever, gastroenteritis, and hepatitis are more common in areas close to waste sites; as this environment favors the breeding and proliferation gastrointestinal infections, flies, and mosquitoes (Harvey et al., 2002; Ziraba et al., 2016; ACCP, 2019). According to WHO (2017), 23% of vector-borne, diarrheal, and cardiovascular disorders in Africa are due to environmental factors and variabilities. And this is supported by Parham et al. (2015), which reported that aside social and demographic factors, environmental factors are one of the major contributors influencing the distribution of vector-borne illnesses. Thus, in addressing this waste-and pollution-driven health and environmental challenges, there is a move for a more sustainable waste management (Abraham *et al*., 2023). The move for a more sustainable waste management and valorization technique opens up new opportunities for innovation, job creation, and the development of a bio-based economy. Waste valorization is any processing activity and technique that aims to reuse, recycle, and/or compost wastes into usable products or energy (Kabongo, 2013). This all-encompassing and progressive strategy emphasizes the potential to turn waste into a resource, fostering a more resilient and sustainable future. Waste valorization has the potential to completely transform how waste is managed worldwide. And with Africa being one of the waste dump hubs with 38% contribution of largest landfills, the opportunities for bio-based economy through waste valorization is enormous. Africa produces about 58% organic waste, and its decomposition contributes significantly to global warming. But this waste can be recycled and converted into valuable products. Microorganisms with their complex biochemical and enzymes play a significant role in waste valorization. They have been implicated in many waste valorization technologies such as in composting, anaerobic digestion, fermentations, etc., to produce biofertilizer, bioenergy, and other value added products. Kavya et al. (2024) highlighted that the use of microbial enzymes’ catalytic properties to transform complicated organic compounds into a variety of valuable products, such as phenolics, flavonoids, anthocyanins, ethanol, bio-oil, essential oils, and nutritional items, is one of the most crucial strategies in microbial-driven waste biomass valorization. In 2018, during the first Africa Waste Management Outlook, Africa was reported to be underperforming with only about 4% recycling of its waste as against the 50% waste recycling projection by 2023 (UNEP, 2018). This might probably be due to the level of interest on waste valorization in this region especially with the use of microorganisms. Thus, the probable missing link between what has been done in terms of research in waste biomass valorization via the use of microorganisms and the hotspots in microbial-driven waste biomass valorization in the African region using the most cited documents, countries, authors’ collaboration and thematic evolution are the main focus of this review using a bibliometric analysis. Systematic review follows different procedural patterns; however, selecting the most relevant research and/or data, as well as determining the possible risks of bias, data extraction method, and how to do an in-depth synthesis of included studies with possible analysis of the meta-data, are all essential systematic review procedure (Afrifa et al., 2022; Alfadil et al., 2022). Figure S1 on the schematic representation of the major concepts in this work summarizes the idea behind systematic and bibliometric review combinations. The use of the PRISMA flow diagram in systematic review has been explored by researchers in recent times (Pant et al., 2017; Drissi et al. 2021; Afrifa et al., 2022). This model flow highlights all the possible ways of eliminating bias and selecting the relevant data for analysis. Here we are asking the following research questions: RQ1: Which authors, journals, and countries are the most prolific in microbial-driven waste biomass valorization research in Africa, and how is their Network? RQ2: How is the evolutionary trend of research in microbial-driven waste biomass valorization in Africa, and who are the contributors? RQ3: What are the current themes and prospects in microbial-driven waste biomass valorization research in Africa?

## 2.0 Research Methodology and Procedures

This study was undertaken to close the gap in the literature on the extent of research in the valorization of organic waste with the aid of microbes in Africa under the current search for alternative waste management strategy by combining bibliometric (quantitative) and systematic (qualitative) analyses. The methodology adopted for this work is elucidated in Fig. 1a, with evidence of the roadmap for systematic review of literature and bibliometric analysis. This roadmap begins with selecting the criteria for inclusion or exclusion of literatures, followed by choice of database for scientific articles, selection of keywords and designing of strings to be used, systematic literature search based on PRISMA 2020 statement, and result analysis (Bibliometrics) using the biblioshiny in the R studio software for the specific database.

**Figure 1:**
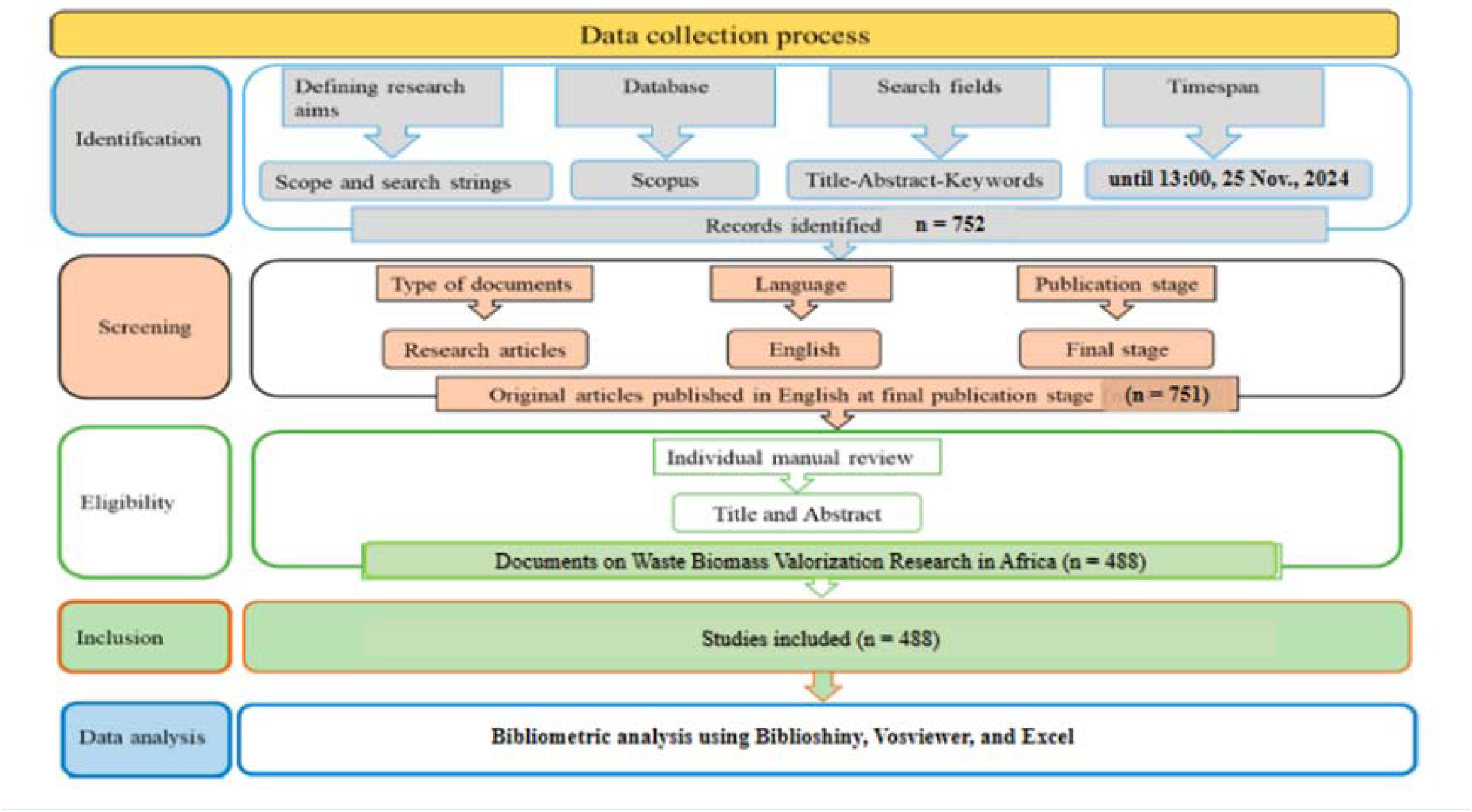
Summary of methodology used for this study.

### 2.1 Database screening

Four well-known and established databases namely: Scopus, ScienceDirect, Dimensions.ai, and Lens.org; chosen mainly due to their accessibility and scope of coverage were first considered for use. According to research Scopus offers access to a vast range of peer-reviewed documents from practically every scientific topic (Al Ryalat et al., 2019; Pranckute, 2021). ScienceDirect contains transdisciplinary publications like Scopus, with a strong emphasis on waste valorization, thus making this database compliant with this study. Dimensions.ai was also selected as it covers a large number of documents and allows for easy downloading and analysis of datasets (García-Sánchez et al., 2019). Len.org was picked due to easy access and the free access to peer-reviewed articles available in it (Rupasinghe et al., 2023). The databases were screened based on the database with the first mention of the word ‘valorization’ and with retrievable data. The selection criteria for using Scopus database in this study highlighted in Table 1.

**Table 1.**
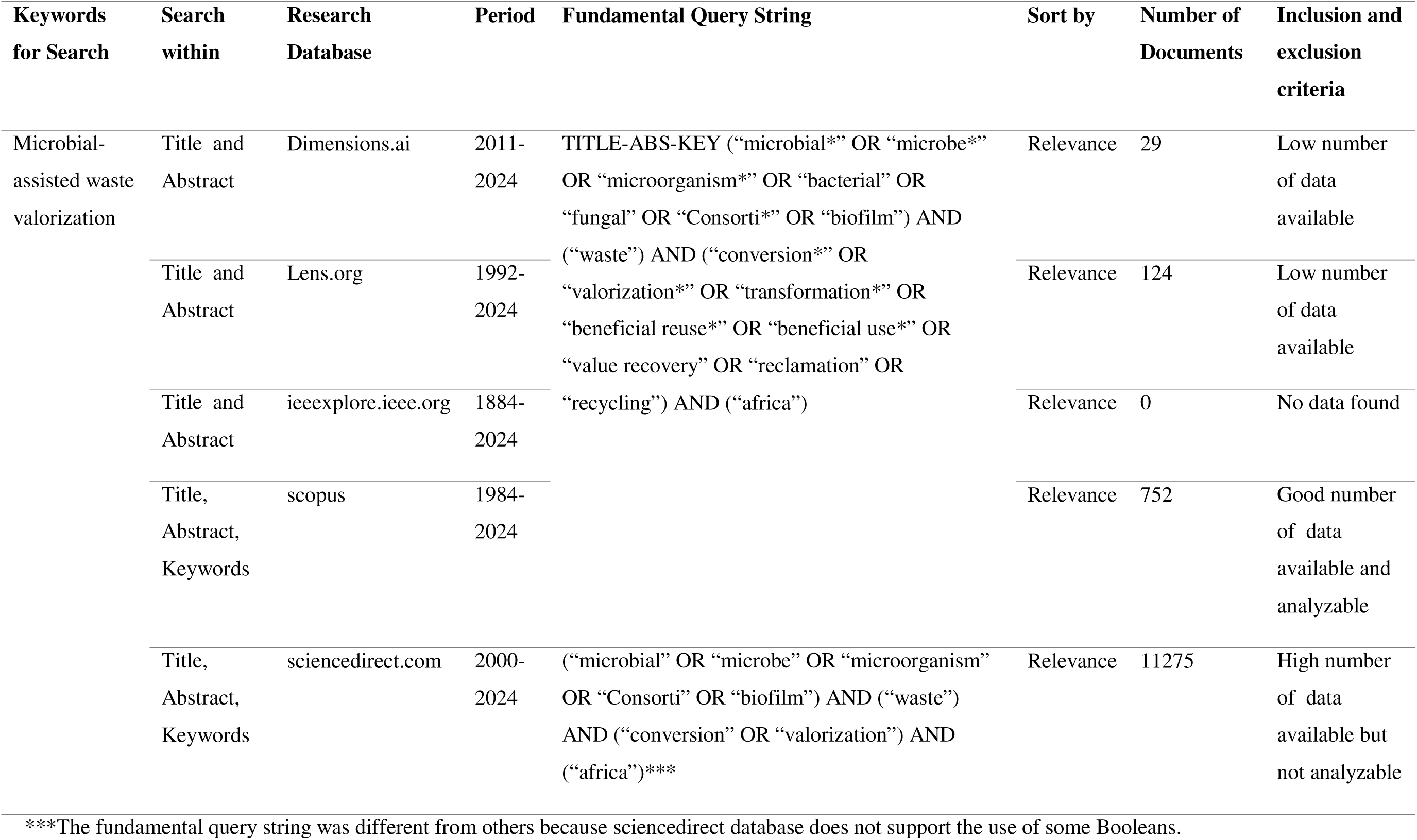
Proprieties and procedures for searching and selection of relevant Database.

### 2.2. Literature Search

A robust search of literature to find out relevant keywords and subjects was done to provide the primary and expert understanding needed for this review and to reduce researcher bias. Thereafter, literature search in scopus database to find out about the role of microorganisms in waste valorization within the African continent, focusing on studies published before November 25, 2024 was done. The criteria for literature inclusion and selection are elucidated in Fig 1b The following terms: ‘microorgamisms’, ‘microbes’, ‘consortia’, ‘biofilm’, ‘waste’, ‘valorization’, ‘recycling’, ‘recovery’ and transformation were considered. However, the search strategy that was eventually used included ((“microbial*” OR “microbe*” OR “microorganism*” OR “bacterial” OR “fungal” OR “Consorti*” OR “biofilm”) AND (“waste”) AND (“conversion*” OR “valorization*” OR “transformation*” OR “beneficial reuse*” OR “beneficial use*” OR “value recovery” OR “reclamation” OR “recycling”) AND (“africa”)).

### 2.3 Study selection

PRISMA model for systematic literature review (SLR) was used to allow for the selection of relevant literatures. The use of the PRISMA flow diagram in systematic review has been explored by researchers in recent times (Pant et al., 2017; Drissi et al. 2021; Afrifa et al., 2022). This model flow highlights all the possible ways of eliminating bias and selecting the relevant data for analysis. This model highlights the identified number of studies, the number of duplicated studies eliminated, the reviewed studies for eligibility and the final included studies. The literature chosen for this study was based on a careful assessment of the abstracts and titles found in the first search results to see if they meet the predetermined inclusion criteria. When necessary, a thorough examination of the complete texts is conducted to determine the admissibility of entries that satisfy these requirements. To guarantee the legitimacy and integrity of the literature chosen for the review, this methodical procedure only uses expert judgment and manual analysis, not any software or artificial intelligence (AI) techniques. Using the conceptual scaffold of PRISMA as basis (Figure 1b), a search string for search in the title, abstract, and keywords of literature in the selected online Scopus database was developed.

**Figure 1b:**
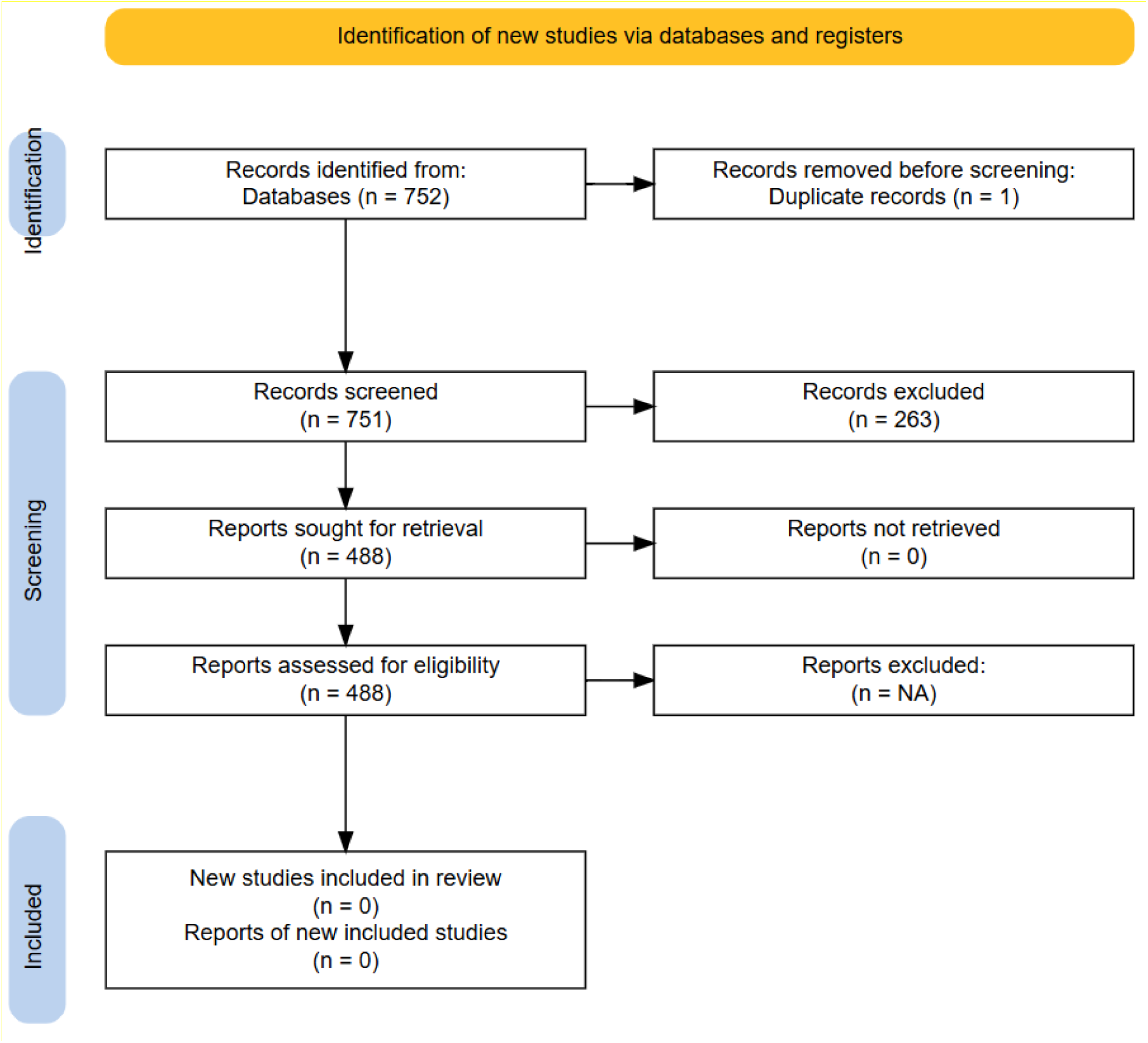
PRISMA flow diagram used for this study.

### 2.4 Inclusion and exclusion criteria

In this systematic review, we focused only on literature that refer to microbial driven waste valorization in situ and ex situ. We removed papers without an abstract and restricted our selection to English-language publications in order to guarantee the specificity and relevance of our findings.

### 2.5 Extraction of qualitative data

Through a rigorous review process, we identified studies that accurately represented the involvement of microbes in waste valorization. From these selected studies, we carefully extracted critical information, such as authorship, year of publication, number of citations, and key findings, thereby providing a robust framework for our analysis. All relevant literature was carefully double checked in accordance with the established inclusion criteria to ensure comprehensive coverage and to prevent any omissions when disparities were observed.

### 2.6 Visual Analysis and Reporting

In order to perform a variety of analyses on the included literature, such as co-authorship network analysis, co-occurrence analysis, and thematic word analysis, we used data visualization and analysis software such as VOSviewer and Biblioshiny. Bibliometrix provided the ability to illustrate scientific trends and productivity within the documents, identifying the most prolific authors and important articles published on the subject; this package includes powerful and comprehensive capabilities for bibliometric analysis, including analyses of authors, institutions, countries, and regions, as well as journal clustering and temporal trends. On the other hand, VOSviewer provides an easy-to-use method of knowledge analysis within research fields, supporting a variety of visualization views to facilitate a deeper understanding of research trends and theme development.

## 3.0 Results

### 3.1 Overview of the analyzed data

Figure 2 denotes the main information on African related microbial-driven waste biomass valorization research performance statistics. The results revealed that microbial-driven waste biomass valorization research in Africa has made significant contributions in the sustainable development goal (SDG) number 11 (sustainable cities and communities) SDG 12 (responsible production and consumption) from 1984 to 2024, judging by the depth of authors (2061) that have contributed in this field, the average citation per documents (25.58) and the volume of relevant document (488) on the subject matter. Figure 3 illustrates the diversity of documents that contributed to this high average citation and annual growth rate; with most of them (66.0%) emanating from Articles, Reviews (22.0%), Book chapters (8.0%), and Conference paper presentations (3.0%). highlights the extent and impact of scientists in the research in microbial-driven waste valorization especially with regards to Africa.

**Figure 2:**
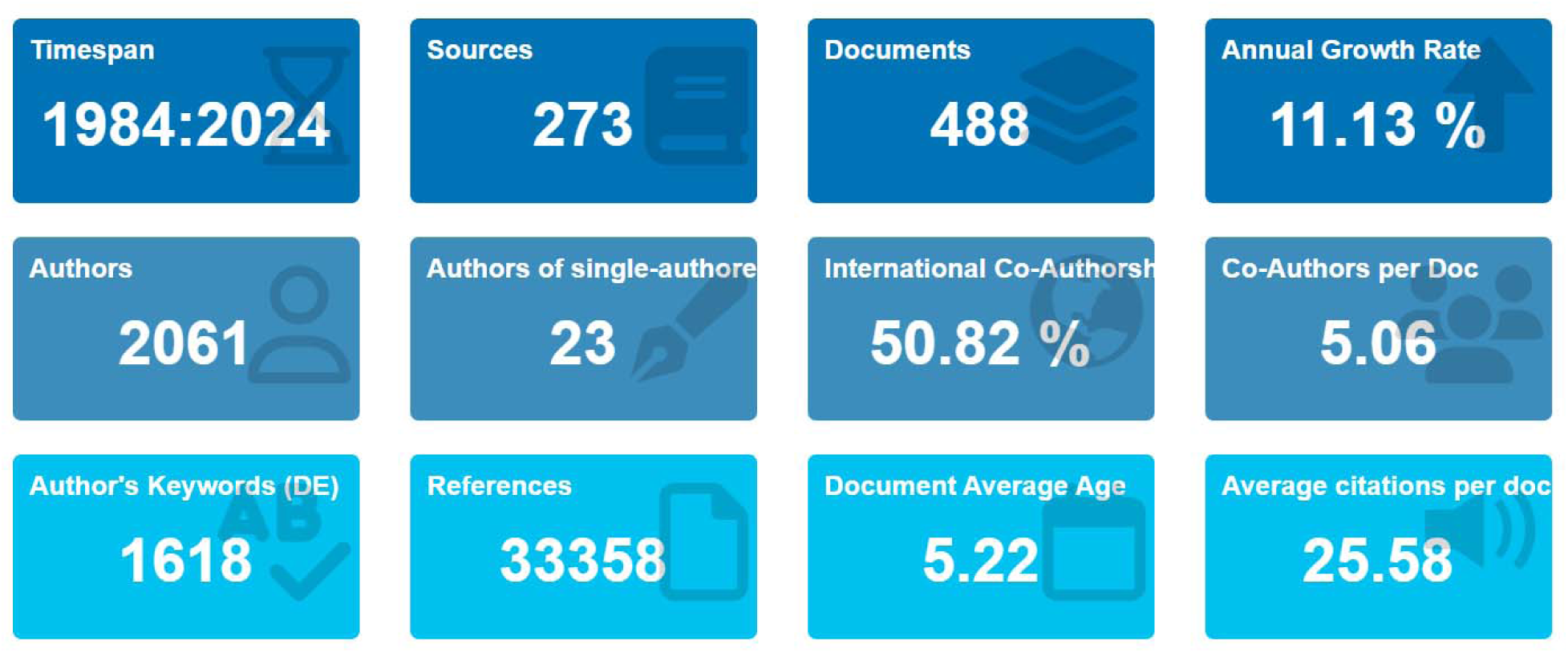
Main Information of performance statistics of microbial-driven waste biomass valorization research in Africa from 1984-2024.

**Figure 3:**
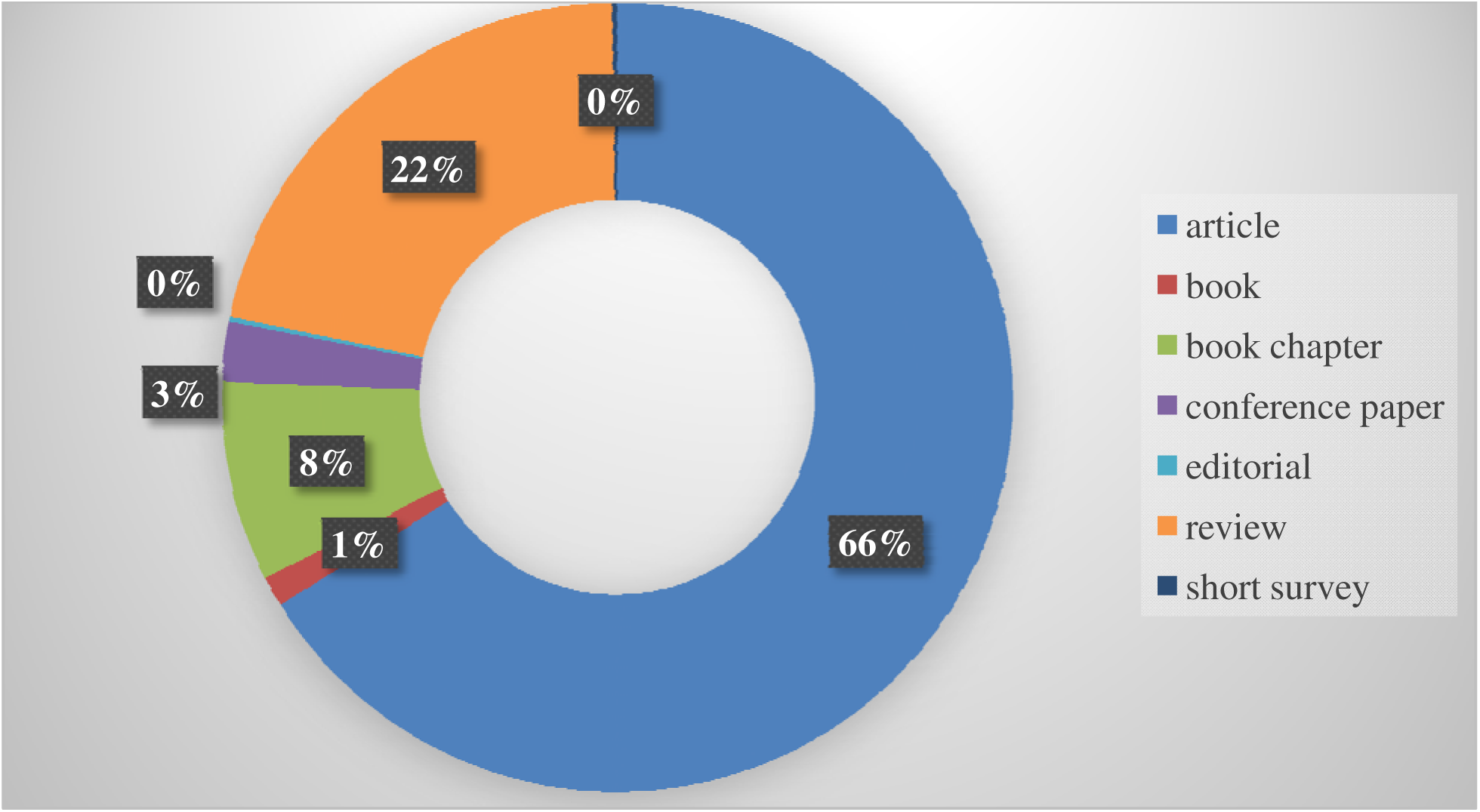
Types of document distribution on microbial-driven waste biomass valorization research in Africa.

### 3.2 Publication and citation per year

The number of documents produced from 1984 to 2024 is denoted in Figure 4. There was a consistent increase in publication around the subject of microbes in waste valorization from 2016 to 2023, and this period is represented as the period of rapid growth. The number of publications on microbial-driven waste biomass valorization with regards to Africa were observed to be redundant and irregular from 1984 to 2005; with most of the years (7 years) having no single publication. However, from 2006 to 2015, there was an observable moderate increase in the number of publications aside 4 documents published in both 2007 and 2012 years. The average yearly publications of 2.72% was observed when the redundant and moderate growth periods (1984-2015) were combined together. On the other hand, the rapid growth period (from 2016 till date) showed a steady rise in the number of publications around the subject of microbial-driven waste biomass valorization in Africa. The average yearly publication of 44.44% was shared between the rapid growth period. The highest number of publications (78 publications) was recorded in 2023. The average yearly citations of documents on microbial-driven waste biomass valorization in Africa is exhibited in figure 5. There was also a gradual fluctuating rise in the number of citations over the period when the research progress was measured based on the yearly mean citation. The highest yearly mean citation of 13.97 was recorded in 2005, followed by year 2021 and 2020 with 7.46 and 7.38 yearly mean citations.

**Figure 4:**
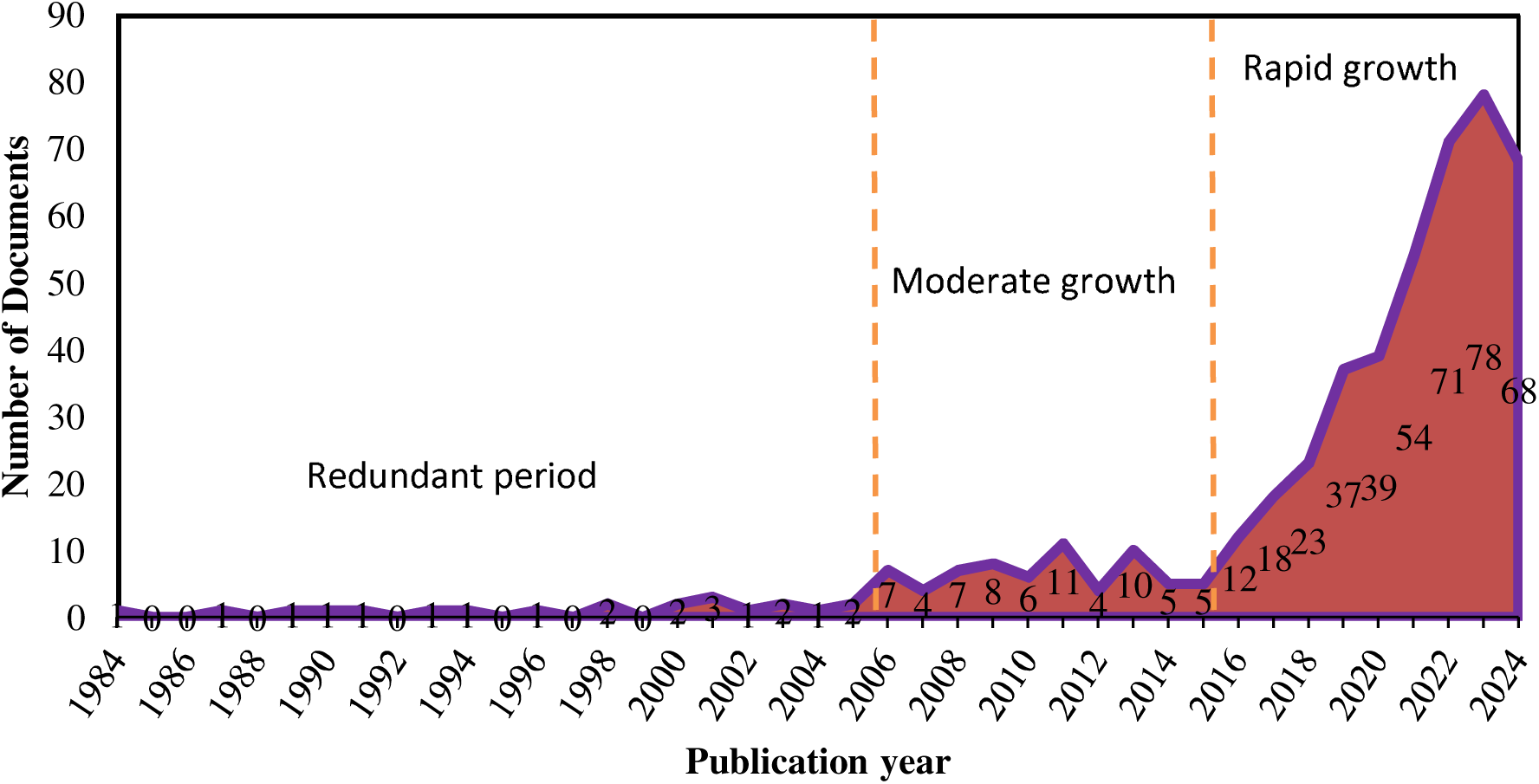
Annual Publication Production on Microbial-driven Waste Valorization Research in Africa.

**Figure 5:**
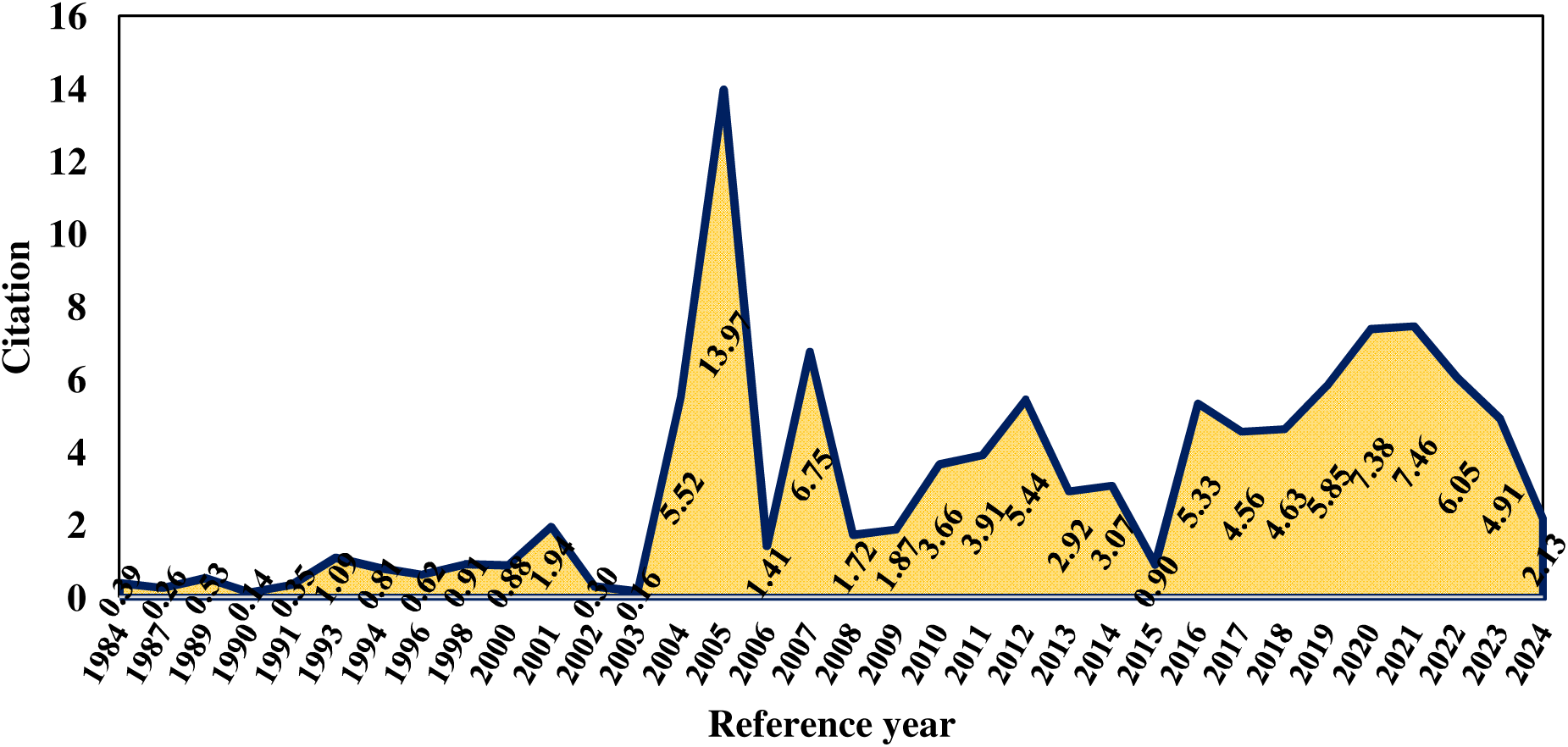
Yearly Mean Citations of Research Publications on Microbial-driven Waste Valorization in Africa.

### 3.3 Publications, citation and corresponding author’s collaborative network by country (top 20)

Figure 6 elucidates the impact of top 20 countries in microbial-driven waste biomass valorization research in Africa based on the number of documents (Fig. 6A) and volume of citations (Fig. 6B). Nigeria, South Africa and Tunisia were ranked as the top 3 countries with 369, 351 and 282 publications respectively; whereas, based on the volume of citations from the publications emanating from these countries, South Africa, Tunisia, and China ranked the top 3 with 3233, 1543, and 1176 citations respectively.

**Figure 6:**
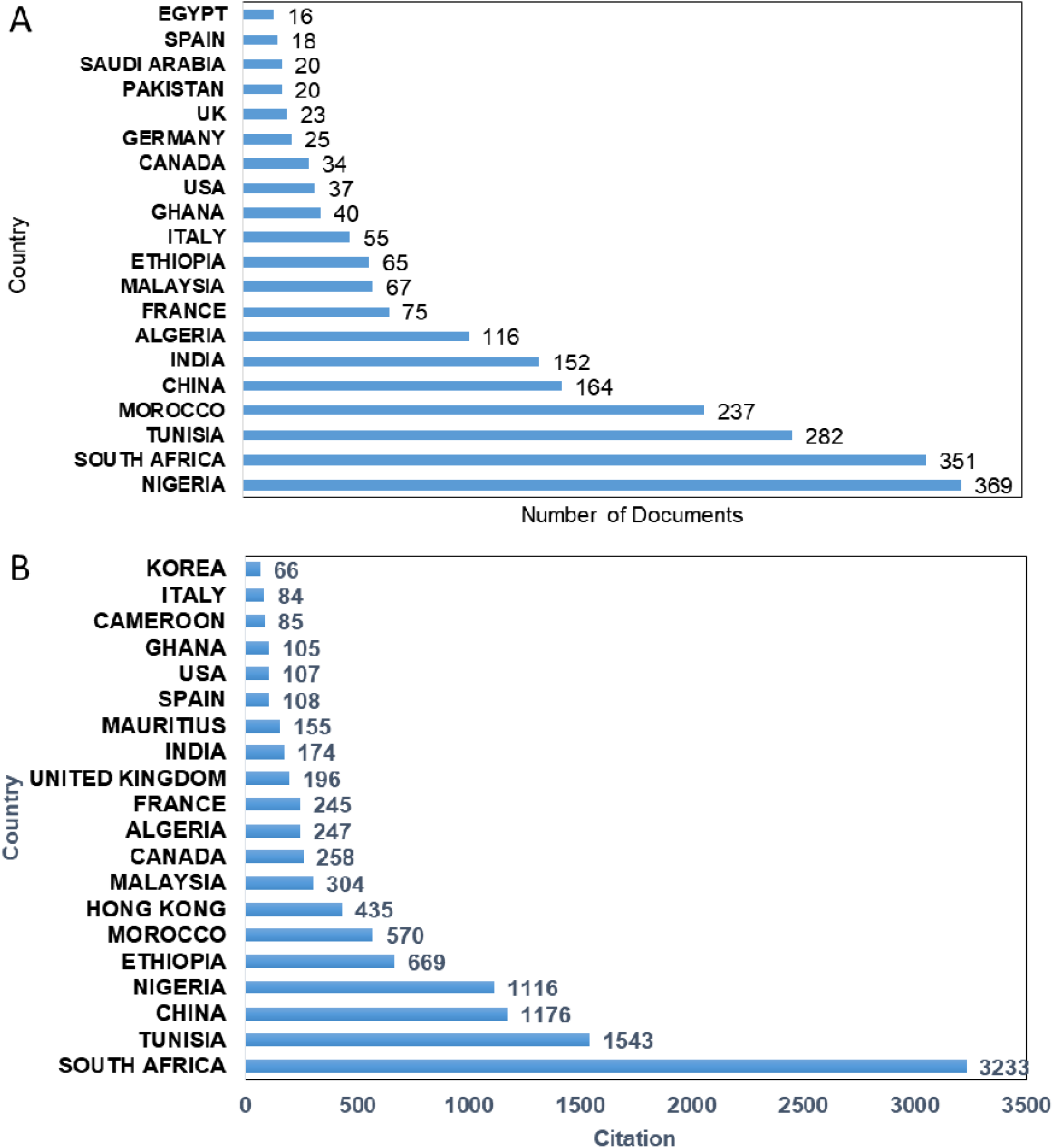
Top 20 Countries Impact in microbial-driven waste Biomass Valorization Research in Africa (A. Number of Publications by Countries; B. Number of Citations)

Figure 7 shows the top 20 countries of document origin based on the corresponding author’s country. South Africa was identified as the country with the highest number of corresponding authors with up to 80 documents coauthored (20 co-authored from multiple countries, and 60 co-authored from a single country). Table 2 highlights the top 20 countries and their collaborations in the subject of microbial-driven waste valorization in Africa. South Africa, Nigeria and Tunisia were ranked top 3 with 60, 43, and 35 collaborations from within the corresponding author country, and 20, 18 and 12 collaborations from outside the corresponding author country, from 80, 61 and 47 articles respectively. Figure 8 depicts the country’scollaboration network on microbial-driven waste biomass valorization in Africa. In figure 8a, South Africa denoted with the biggest light blue circle (cluster 1) boasted up to 30 inter–country collaborations, with the collaborations mostly with countries like India (with 10 documents shared between them), followed by China (8), and Italy (8). In cluster 2, Nigeria, denoted with the biggest dark red circle, had 22 inter-country collaborations; however, India, South Africa, and Malaysia with 12, 12 and 11 documents respectively shared with Nigeria, were countries with the most inter-country collaboration. Cluster 3 (deep purple circles) depicts India leading the cluster with up to 34 inter countries collaborations.

**Figure 7:**
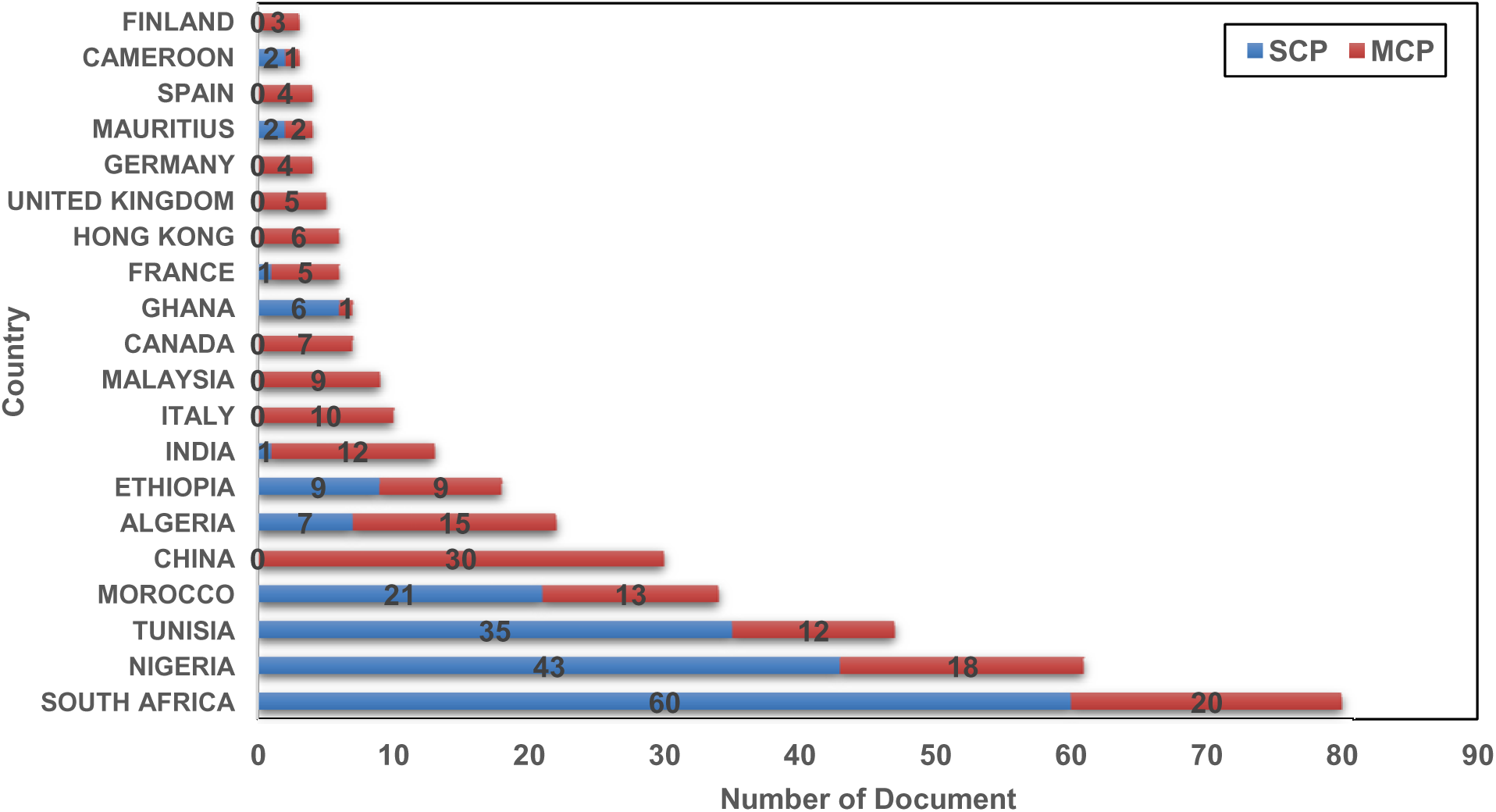
Corresponding Author’s Countries SCP (Single Country Publication or Intra-country collaborations) and MCP (Multiple Countries Publication or Inter-country collaborations)

**Figure 8:**
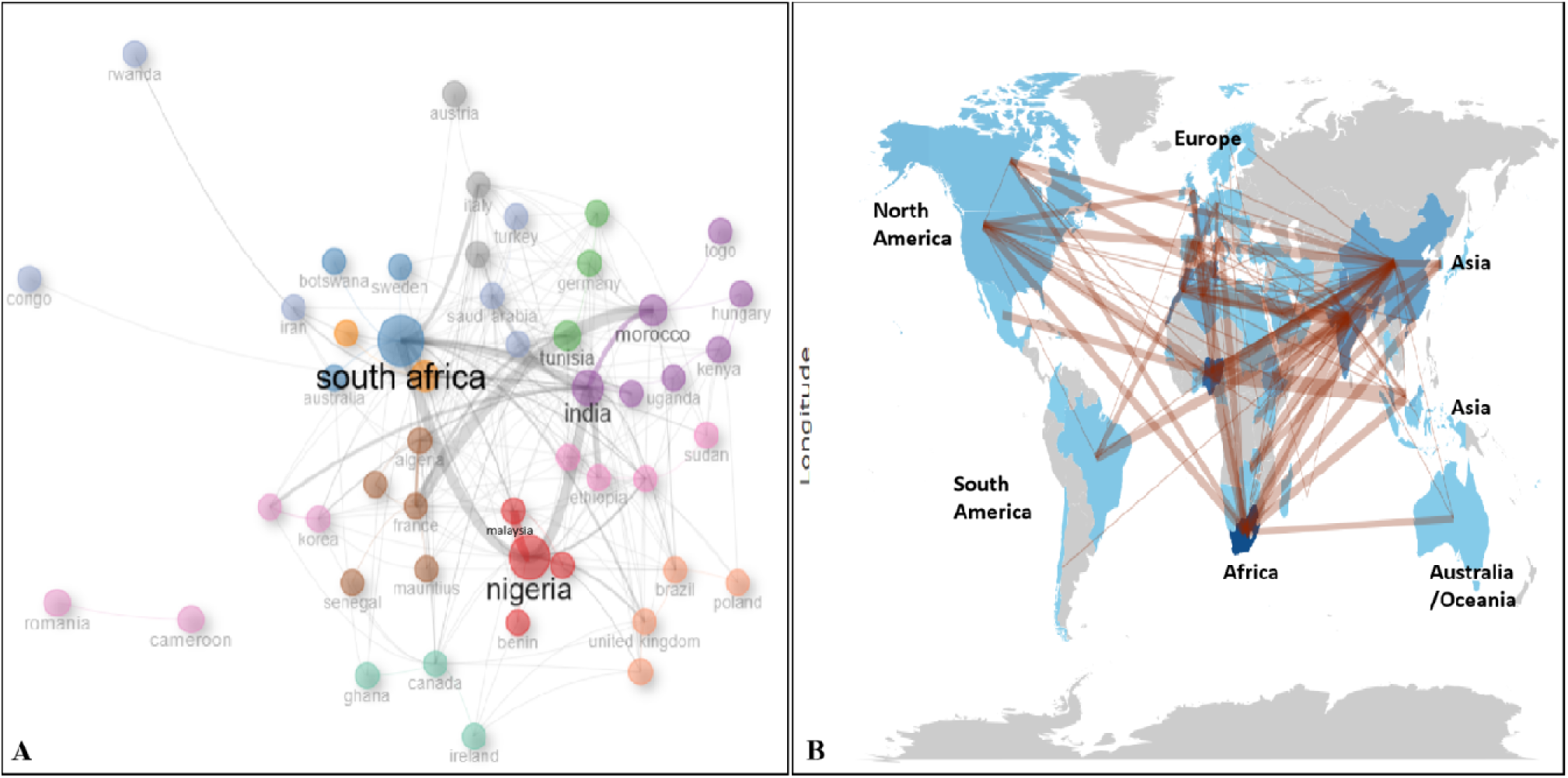
Country’s Collaboration Network on Microbial-driven waste Biomas Valorization in Africa (A: Inter-country collaboration; B: Intercontinental collaboration)

**Table 2:** Collaboration Distribution Based on Corresponding Author Country (Top 20)

| Country | Articles | Articles % | SCP | MCP | SCP% | MCP % |
| --- | --- | --- | --- | --- | --- | --- |
| **SOUTH AFRICA** | 80 | 16.4 | 60 | 20 | 75.0 | 25 |
| **NIGERIA** | 61 | 12.5 | 43 | 18 | 70.5 | 29.5 |
| **TUNISIA** | 47 | 9.6 | 35 | 12 | 74.5 | 25.5 |
| **MOROCCO** | 34 | 7 | 21 | 13 | 61.8 | 38.2 |
| **CHINA** | 30 | 6.1 | 0 | 30 | 0.0 | 100 |
| **ALGERIA** | 22 | 4.5 | 7 | 15 | 31.8 | 68.2 |
| **ETHIOPIA** | 18 | 3.7 | 9 | 9 | 50.0 | 50 |
| **INDIA** | 13 | 2.7 | 1 | 12 | 7.7 | 92.3 |
| **ITALY** | 10 | 2 | 0 | 10 | 0.0 | 100 |
| **MALAYSIA** | 9 | 1.8 | 0 | 9 | 0.0 | 100 |
| CANADA | 7 | 1.4 | 0 | 7 | 0.0 | 100 |
| GHANA | 7 | 1.4 | 6 | 1 | 85.7 | 14.3 |
| FRANCE | 6 | 1.2 | 1 | 5 | 16.7 | 83.3 |
| HONG KONG | 6 | 1.2 | 0 | 6 | 0.0 | 100 |
| UNITED KINGDOM | 5 | 1 | 0 | 5 | 0.0 | 100 |
| GERMANY | 4 | 0.8 | 0 | 4 | 0.0 | 100 |
| MAURITIUS | 4 | 0.8 | 2 | 2 | 50.0 | 50 |
| SPAIN | 4 | 0.8 | 0 | 4 | 0.0 | 100 |
| CAMEROON | 3 | 0.6 | 2 | 1 | 66.7 | 33.3 |
| FINLAND | 3 | 0.6 | 0 | 3 | 0.0 | 100 |
| Average | 18.7 | 3.8 | 9.4 | 9.3 | 29.5 | 70.5 |

### 3.4 Publication by institutions (top 20)

Figure 9 illustrates the top 20 academic institutions contributing the most to microbial-driven waste biomass valorization in Africa. Durban University of Technology, and University of Fort Hare, all in South Africa, had the highest number of publications, with 59 and 35 documents respectively. Following University of Fort Hare, is University of Nigeria, Nigeria, with 31 publications, Stellenbosch University and University of Pretoria, South Africa (30), University of Sfax, Tunisia (29), University of Carthage, and University of Manouba Tunisia, (24), University of Technology Malaysia (22), Federal University of Technology, Nigeria, (21), Covenant University, Nigeria, and The Hong Kong University of Science and Technology, Hong Kong (20), Donghua University, China (17), Landmark University, and Northwest University, Nigeria, (16), Cadi Ayyad University, Morocco, and National Institute of Applied Science and Technology, Tunisia, (15) and Ladoke Akintola University of Technology, Nigeria, Marrakesh, Morocco, and Universiti Teknologi Petronas (14).

**Figure 9:**
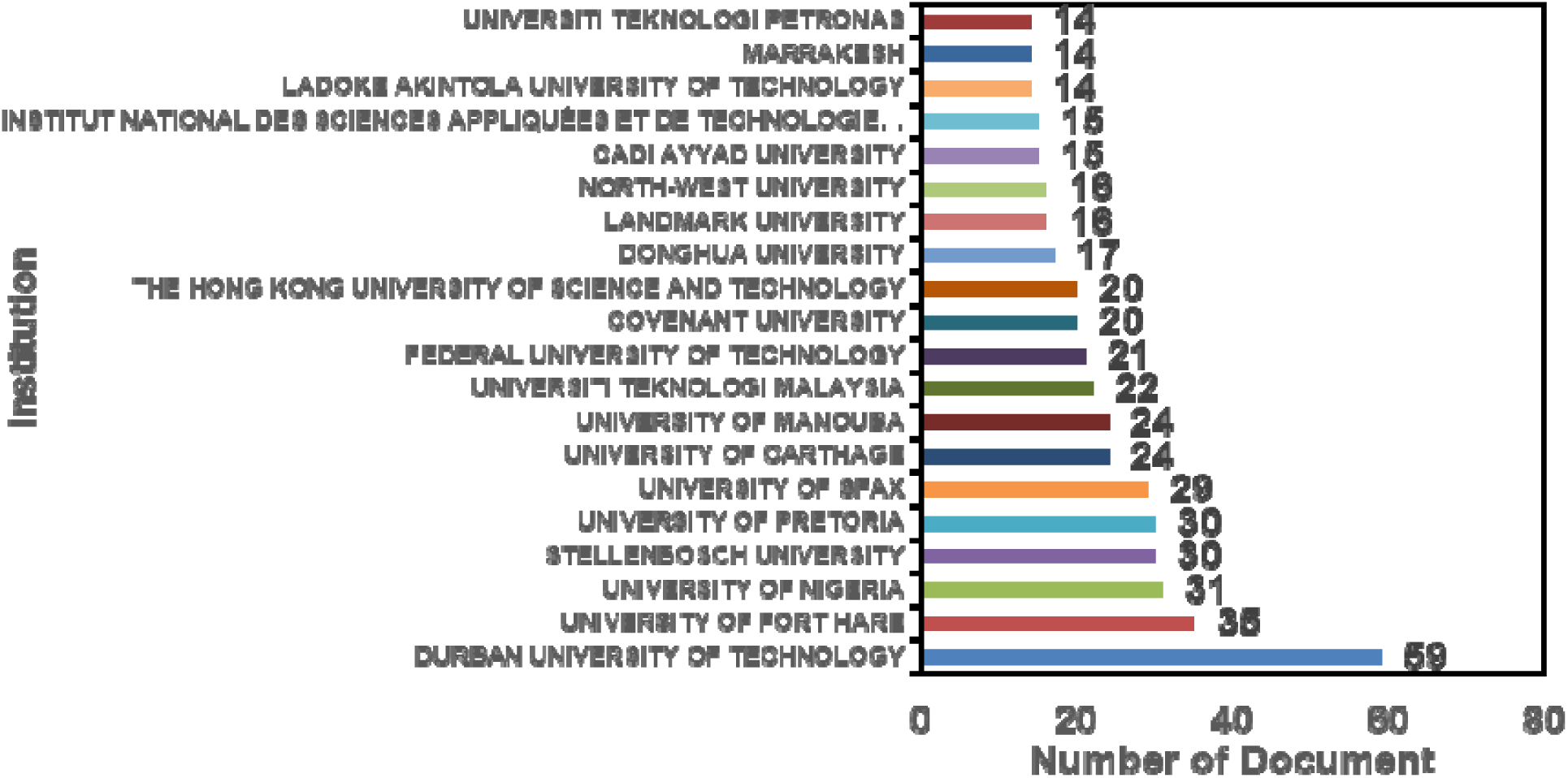
The Leading Institutions on microbial-driven waste Biomass Valorization Research in Africa (Top 20)

### 3.5 Publication by journals (top 20)

The top 20 most relevant journal sources publishing articles on microbially-driven waste biomass valorization in Africa are presented in Figure 10. Generally, it was observed that the top 20 journals published 173 documents, accounting for approximately 35.5% of the total sampled 488 papers. From Figure 10, Bioresource Technology ranks first on the list, with 30 documents corresponding to about 17.3% of the documents published by the top 20 journals. This was followed closely by Journal of Environmental Management, and Science of the Total Environment with 15 published documents each, and Biomass Conversion and Biorefinery with 13 documents.

**Figure 10:**
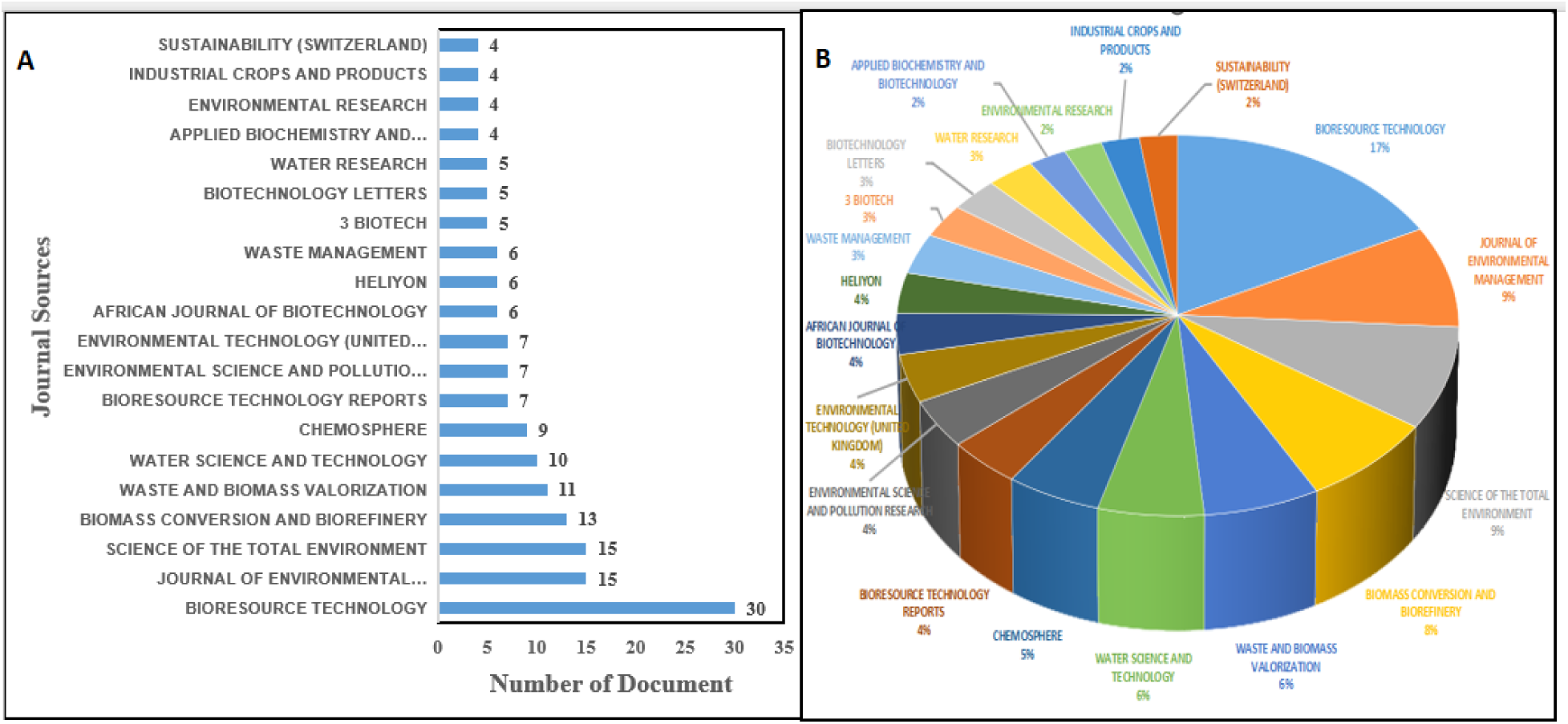
Most Relevant Sources on microbial-driven waste Biomass Valorization Research in Africa; (A) Based on Document Counts (Top 20); (B) Based on h-index Impact Factor (Top 10)

### 3.6 Top 20 leading authors

#### 3.6.1 According to number of publications

Figure 11 depicts the top 20 leading authors in microbially-driven waste biomass valorization research in Africa. From 1944 to 2024, 2,061 authors were involved in microbially-driven waste biomass valorization research in Africa. The findings showed that Hamdi M, Hafidi M, and Bux F were the leading top 3 authors in microbially-driven waste biomass valorization research in Africa with a total of 34 publications which is about 1.4% of the total publications. Hamdi M. is ahead of other 2060 authors as the most prolific author with 15 documents, followed by Hafidi M. with 10 documents, Bux F. with 9, Nwodo UU with 8, and Nnolim and Van Zyl WH with 7 documents each.

**Figure 11:**
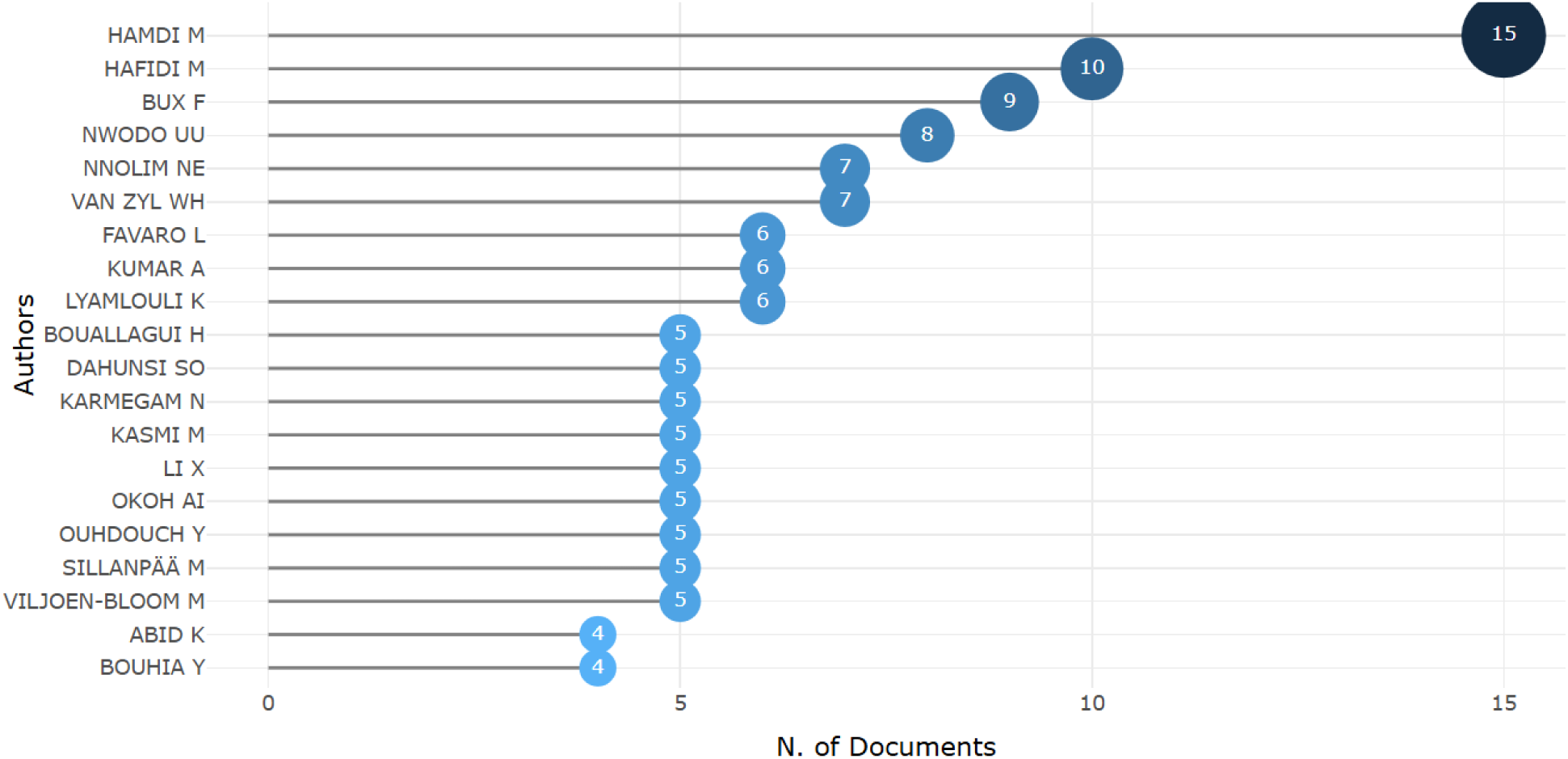
Leading Authors on microbial-driven waste Biomass Valorization Research in Africa (Top 20)

#### 3.6.2 According to active productive years, volume of publications and citations

Figure 12 illustrates a graph depicting the productivity of the top 20 authors over time, quantifying and charting their output in terms of publication count and impact across a specified period. The metrics in this figure 12 presents an overview of the 20 most prolific authors during the past 25 years (2000–2024), with productivity assessed based on the volume of publications by each author during this time-frame and the accompanying citation accrued by the author. The length of line shows the range of author’s activity based on publications, while the size of bubbles shows the volume of documents published and the intensity of color reveals the volume of citation for such author works in a particular year. For instance, Hamdi M had an output and publication productivity (red line) over a 23 year period (2000-2022); with an approximately 1.69 total citations per year in the start year of 2000 for only one document published.

**Figure 12:**
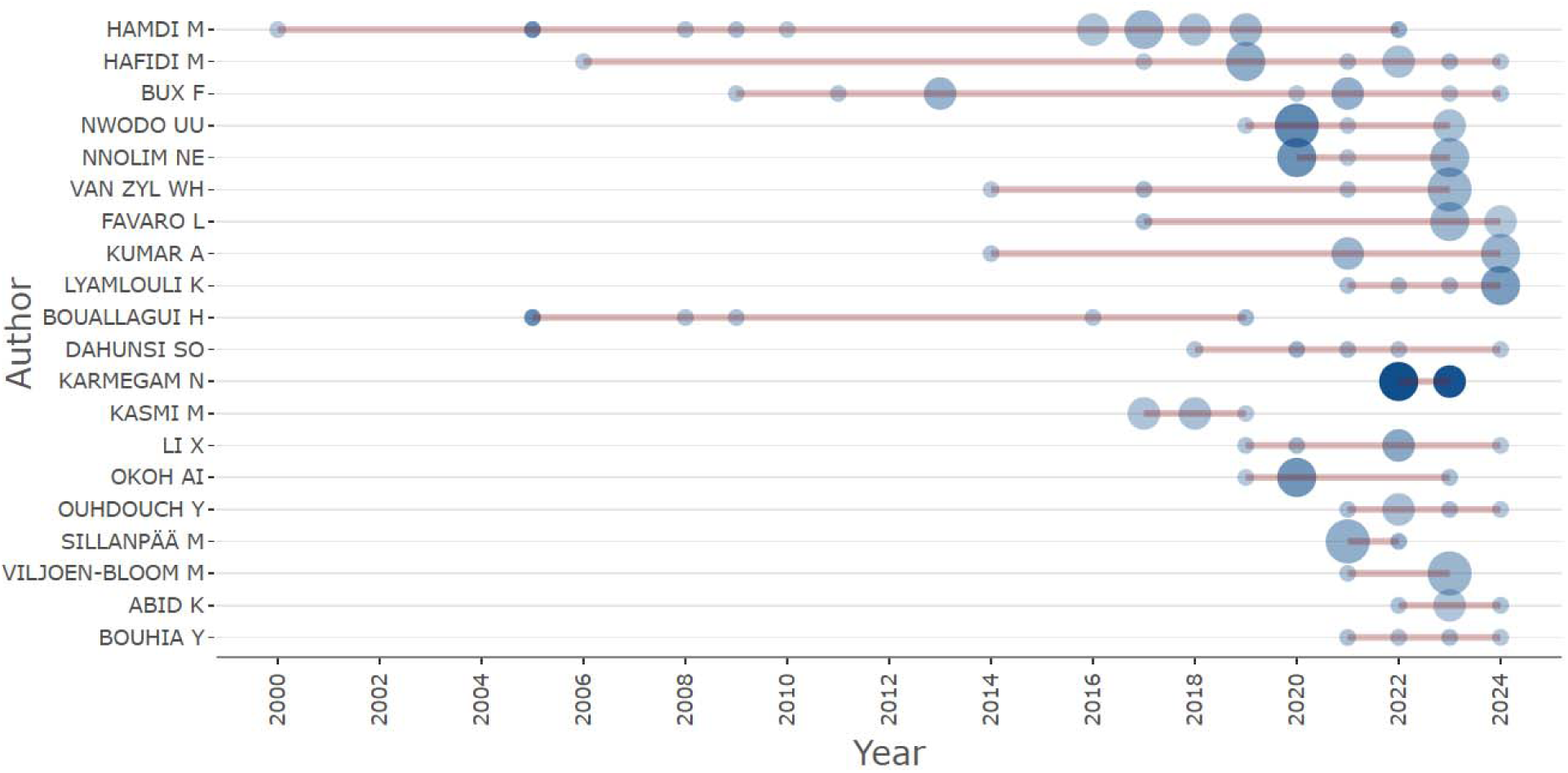
Top 20 Authors’ Productivity on microbial-driven waste Biomass Valorization Research in Africa.

#### 3.6.3 According to h-, g-, and m-indexes

Supplementary Table S1. presents the 20 most significant author impacts arranged by h-index. Hamdi M leads the table with 12, 15, and 1003 h-index, g-index and total citation (TC) respectively, with 0.48 m-index after 25 years. The author with the highest m-index (1.667) is Armegam N with a total citation of 276 after 3 years.

### 3.7 Most-cited documents based on Citations (Top 11)

Table 3 shows the top 11 most cited documents in microbial-driven waste biomass valorization in Africa based on Scopus’s data analyzed, with all these documents spanning across seventeen years (2005 to 2021). Out of the 12483 total citations from the 488 documents analyzed, 25.4% (3168) of the total citations were recorded from the top 11 most cited documents with up to 100 citations. The least and highest citations were recorded in the works by Nsenga et al. (2019) and Bouallagui et al. (2005) with 105 and 557 citations respectively.

**Table 3:**
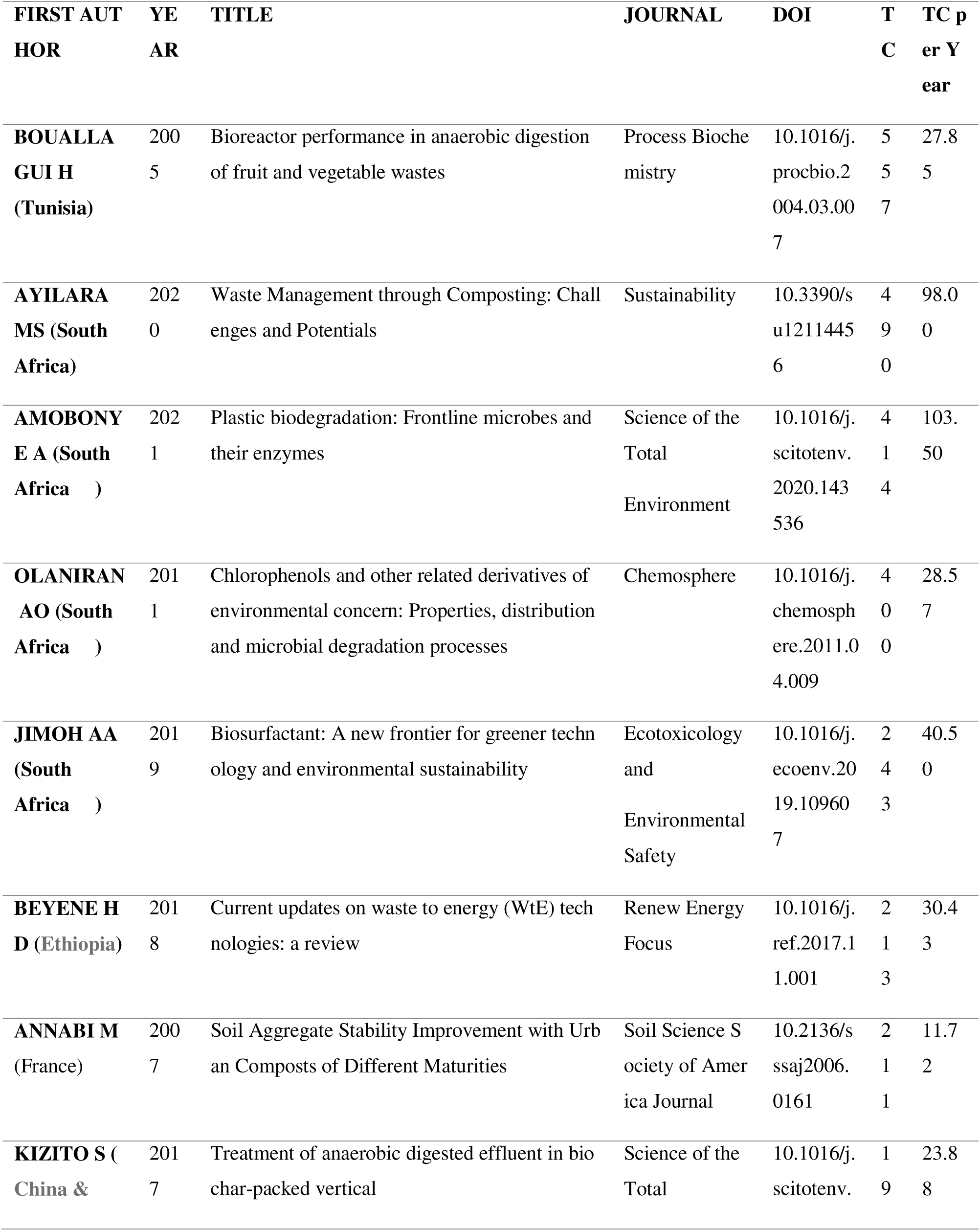
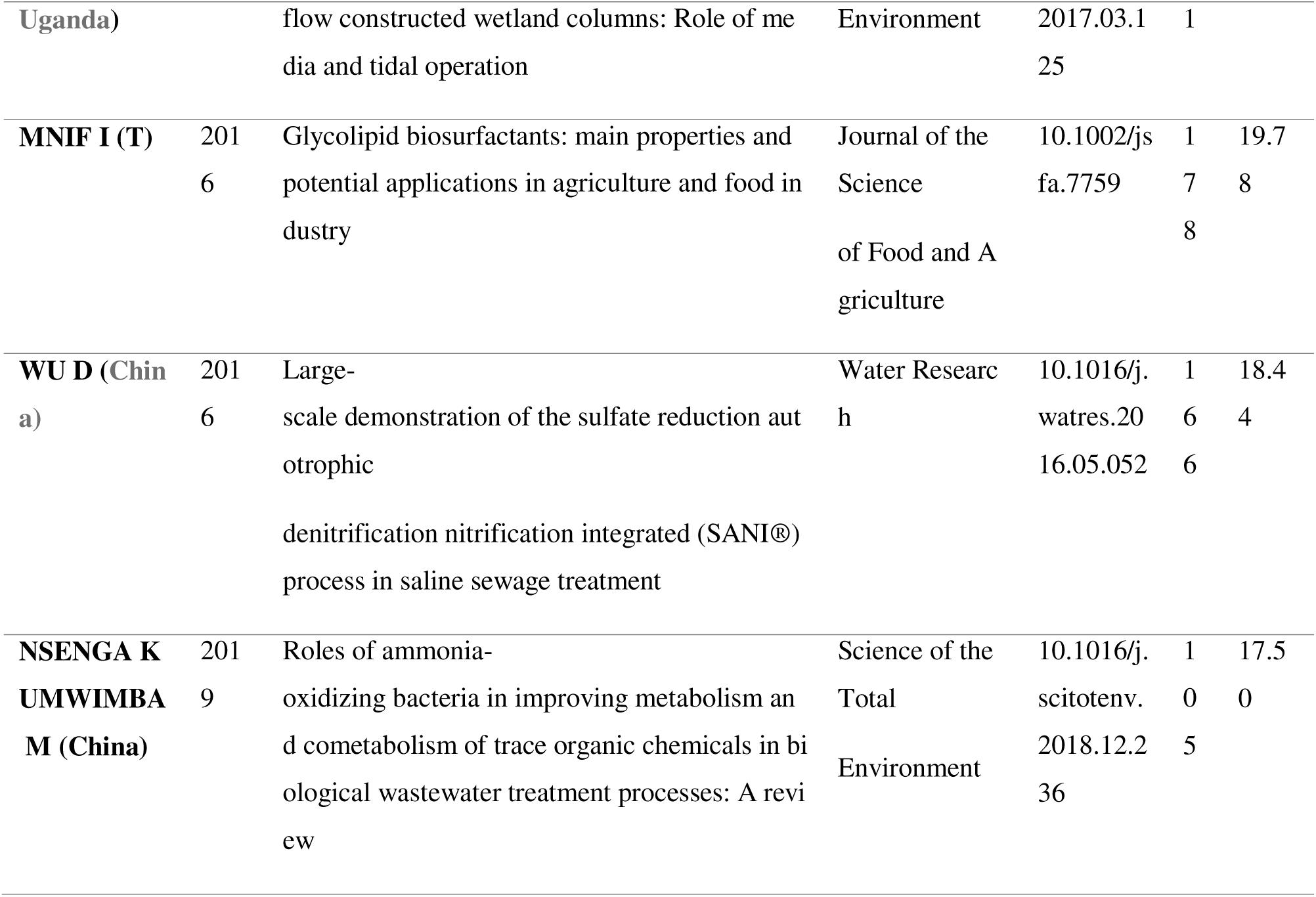
Most Globally Referenced Documents in Research on microbial-driven waste Biomass Valorization in Africa (Top 11)

### 3.8 Topical issues, Emerging Trends and Research Gaps

#### 3.8.1 Trending topics

Figure 13 shows that the evolutionary topical focuses in researches on waste biomass valorization from 2001 to 2024. From 2001 to 2006, the trending topics were about the valorization of molasses from sugarcane to the use of phanerochaete species, and *Enterobacter aerogenes* in waste management. From 2007 to 2012, there seemed to be increased interest in the search for beneficial uses of agro/industrial wastes and soil microbes. Both wastes from the processing of cassava (*Manihot esculenta*) and textile waste water were also another topic discussed within this period (2007 to 2012). From 2013 till 2019, attention began shifting from alcohol production from waste to anaerobic digestion of waste and enzymatic processes in waste management. From 2020 till 2024 when this review was done, the trending issues were narrowly around waste valorization as different concepts were seen speaking to these issues. The trending topics ranged from recycling, wastewater treatment, anaerobic digestion, enzymatic hydrolysis, valorization, plastic recycling, purification etc. (Figure 13).

**Figure 13:**
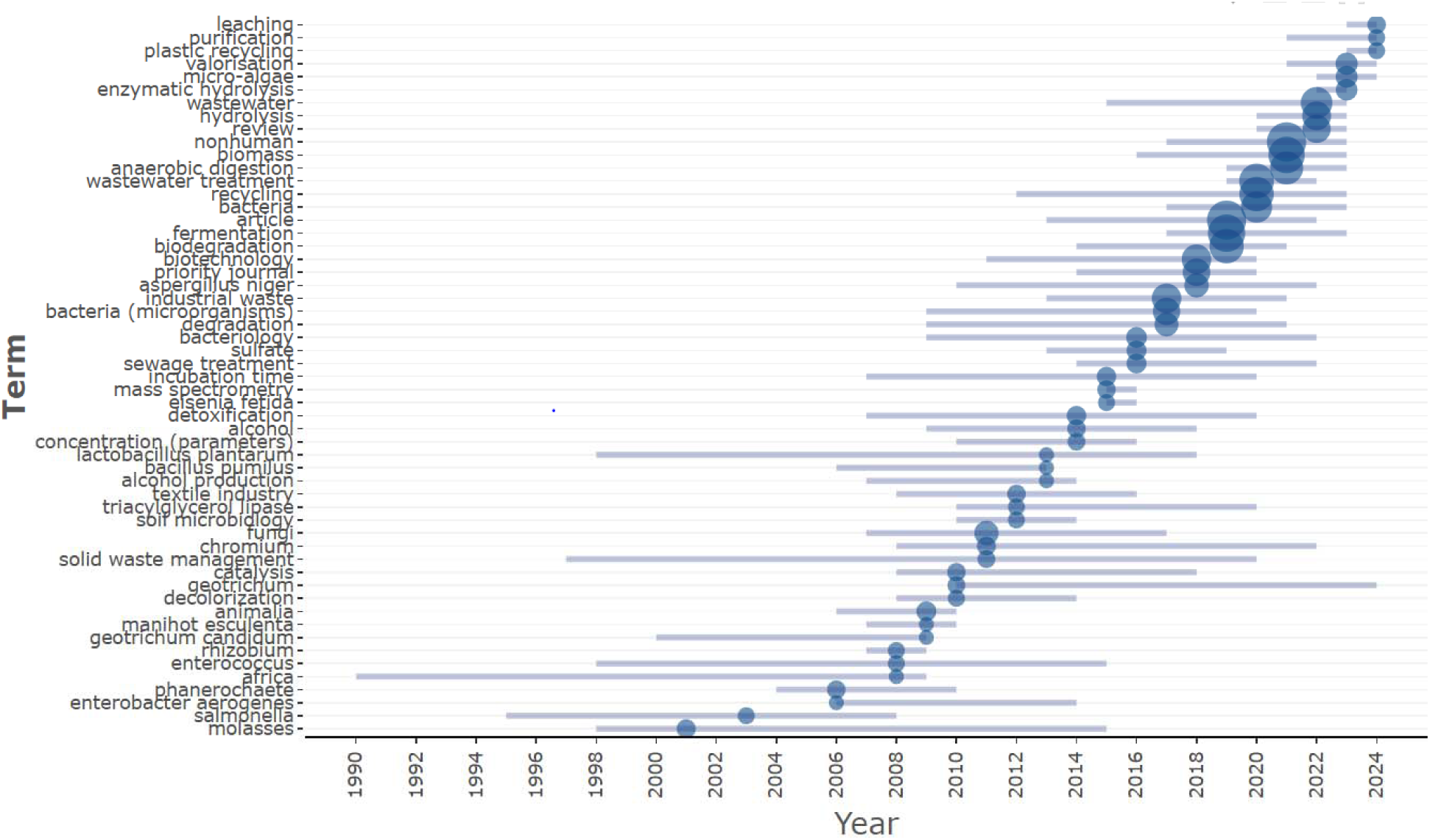
Trending Topics on microbial-driven waste Biomass Valorization Research in Africa.

**Figure 14:**
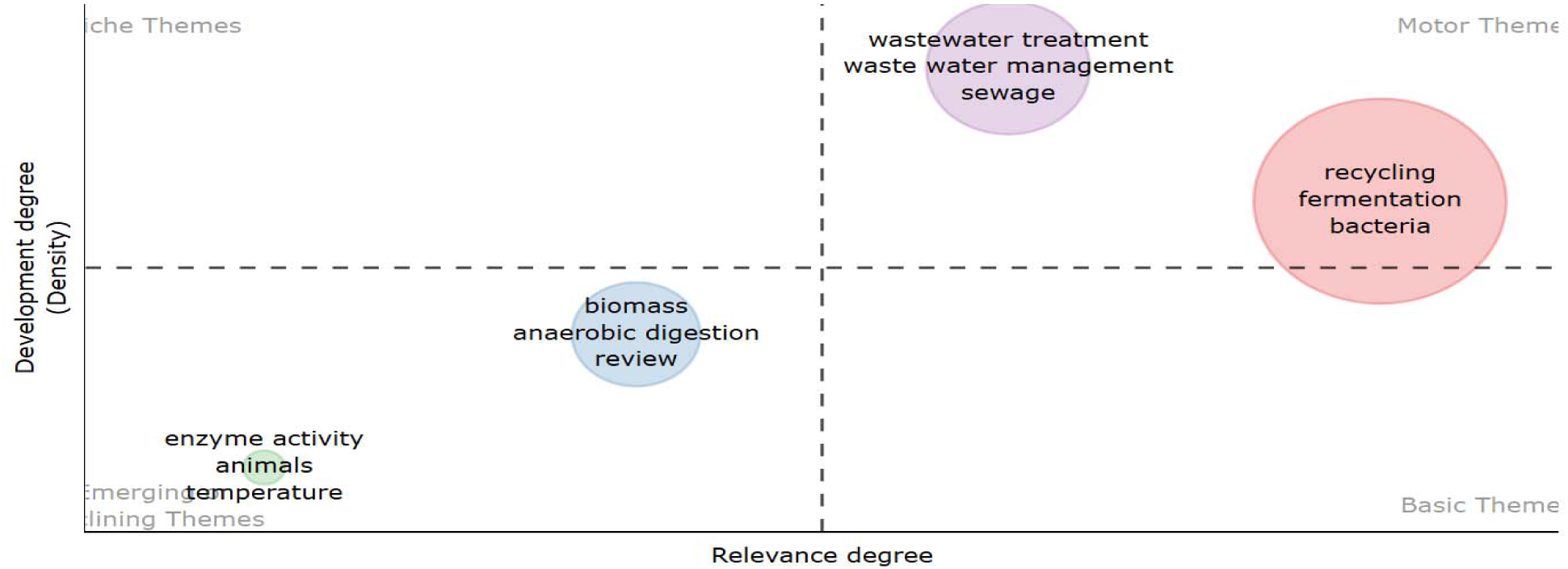
Thematic map from keywords plus on microbial-driven waste Biomass Valorization Research in Africa.

#### 3.8.2 Thematic evolution and trends based keyword plus

The thematic trend map consists of four quadrants depicting the four levels and degree of relevance of these keywords in the past, currently and the future. The lower left quadrant (emerging and declining themes) encompasses subjects characterized by low density and focus, indicative of themes that have recently emerged or vanished. Two clusters are shown in this quadrant based on the color. The light green cluster consisting of enzyme activity, temperature, etc. tends to reveal the declining themes, while the light blue cluster consisting of biomass, anaerobic digestion, etc tends to show emerging themes as corroborated by trending topics and thematic evolution in figure 12 and 14 respectively. The region in the upper left quadrant (Niche themes) denotes a specific topic mostly excluded from the primary emphasis which could be said to be a niche in research on microbial-driven waste biomass valorization. The regions in the upper right quadrant (motor themes) signify well-established and resilient subjects within the domain of research on microbial-driven waste biomass valorization. This quadrant contains two clusters signified with pink and purple colors. The pink color cluster reveals highly relevant themes with slow degree of development; whereas the purple cluster shows themes that are of seldom relevance and high developmental degree. The regions in the lower right quadrant denote the most prevalent fundamental and transverse subjects, however internal development is not as enhanced as that of the upper right quadrant. The themes in the lower left and right quadrants may serve as a source of inspiration and potential reference for more research in microbial-driven waste biomass valorization.

#### 3.8.3 Thematic evolution and trends based keyword plus

Figure 15 denotes the thematic evolution of the keyword plus based on the dataset from 1984 to 2024. Three periods were chosen as cut-off points: 2005, 2016, and 2017 based on the yearly publication and citation results (Figure 4 and 5), where it was observed that there were significant increases in the number of publications and citations across these periods. In the first period (1984–2005), the popular keywords were cellulase, biodegradation, anaerobic digestion, waste and waste water management, which merged into the next period (2006–2016) as biomass, wastewater, and animal. The keywords in the second period metamorphosed into nonhuman, wastewater treatment and anaerobic digestion in the last period (2017-2024).

**Figure 15:**
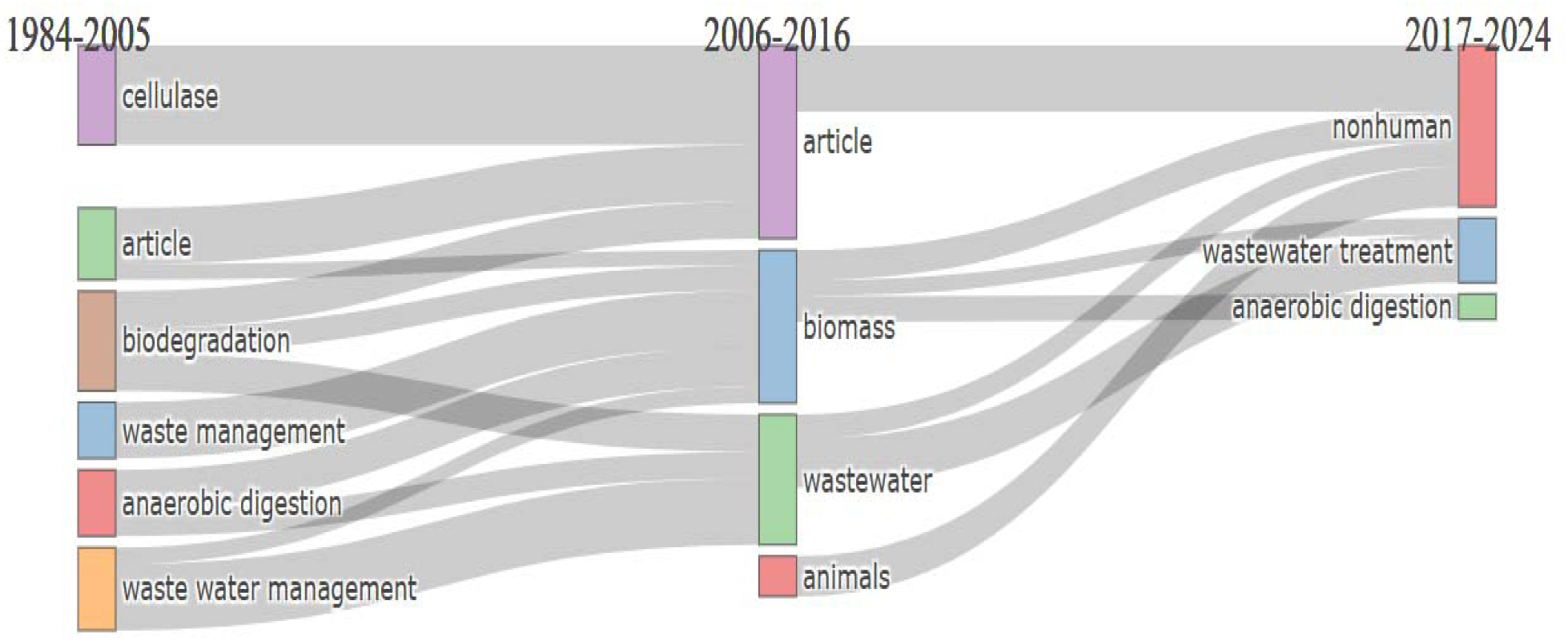
Thematic Evolution of Author Keywords on microbial-driven waste Biomass Valorization Research in Africa.

### 3.9 Keywords co-occurrence analysis

#### 3.9.1 Co-occurrence—all keywords

Keywords are a very significant component of scholarly articles speaking to the most re-occurring words that characterizes a literature; and this plays a crucial role in the type of information retrieved from a publication. Figure 16 shows all keywords co-occurrence network drawn using VosViewer which was used to analyze all keywords re-occurring in microbial-driven waste biomass valorization research in Africa. The keywords are arranged in 7 clusters signified by the frames with different colorations and sizes. Figure 16a showed all author keywords with Network visualization. The cluster 1 represented by the red framed keywords has 135 keywords clustered in it. Cluster 2, represented by green, contains 102 keywords, while Clusters 3 and 4, represented by blue and yellow, have 100 and 99 keywords clustered in them. Finally, cluster 5, 6 and 7, represented by purple, turquoise blue and orange color respectively contain 98, 28 and 12 keywords respectively.

**Figure 16:**
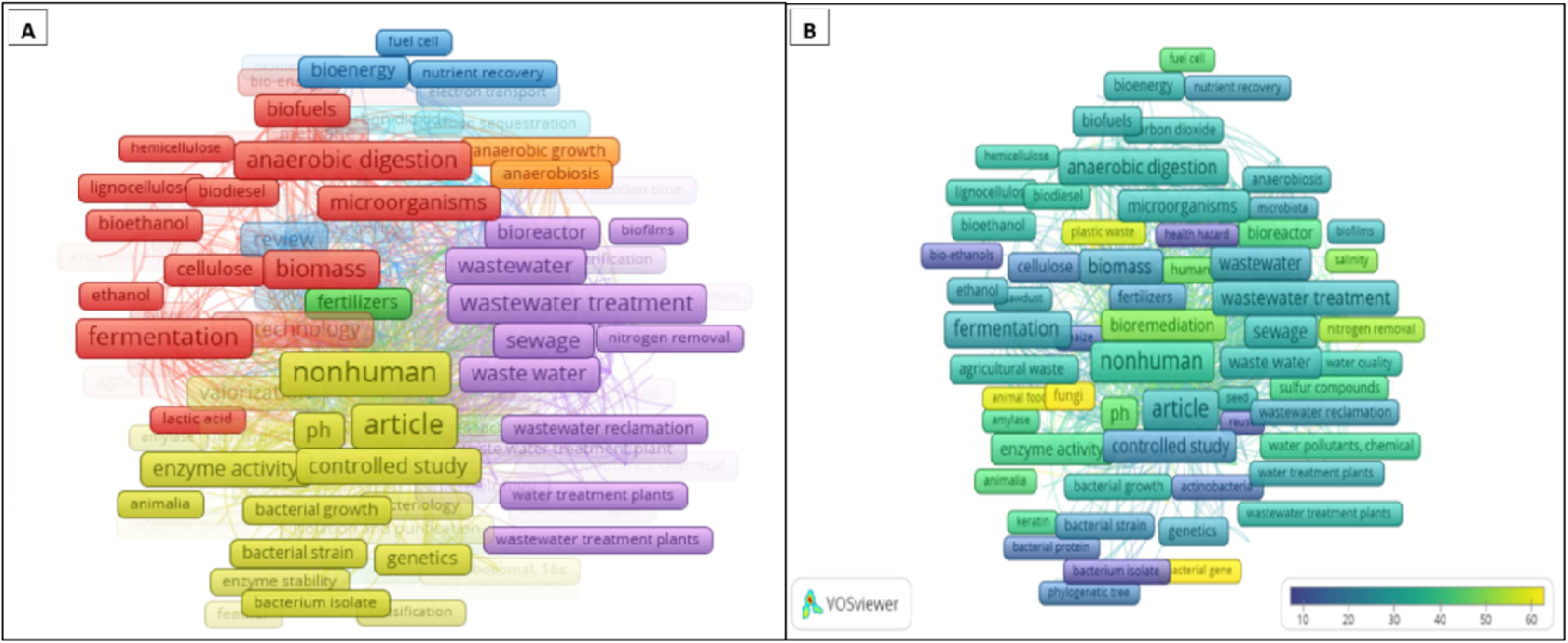
All Keywords Co-occurrence Network (A) and overlay by average yearly citation from microbial-driven waste Biomass Valorization Research in Africa.

#### 3.9.2 Co-occurrence—author keywords

This set of keywords are primarily provided by the author. It is an excerpt from the all keywords. The same minimum keyword occurrence of 5 was selected; and out of the 1614 keywords from the different authors, only 57 met the criterion. Figure 17 depicts author keywords co-occurrence network (fig. 17a) and overlay visualization by average yearly publication (fig. 17b) in microbial-driven waste biomass valorization research in Africa. Eight (8) clusters having a total link strength of 363 and distinct colors are shown in the co-occurrence network. From figure 17a, cluster 1 (red framed keyword), 2 (green framed keyword) and 3 (blue framed keyword) each consist of 10 keywords. The yellow color framed keywords represent cluster 4 (9 keywords). Cluster 5 represented in purple contained 8 keywords, while Clusters 6 (5 keywords), 7 (4 keywords) and 8 (1 keyword), are represented by torques blue, orange and brown colors respectively.

**Figure 17:**
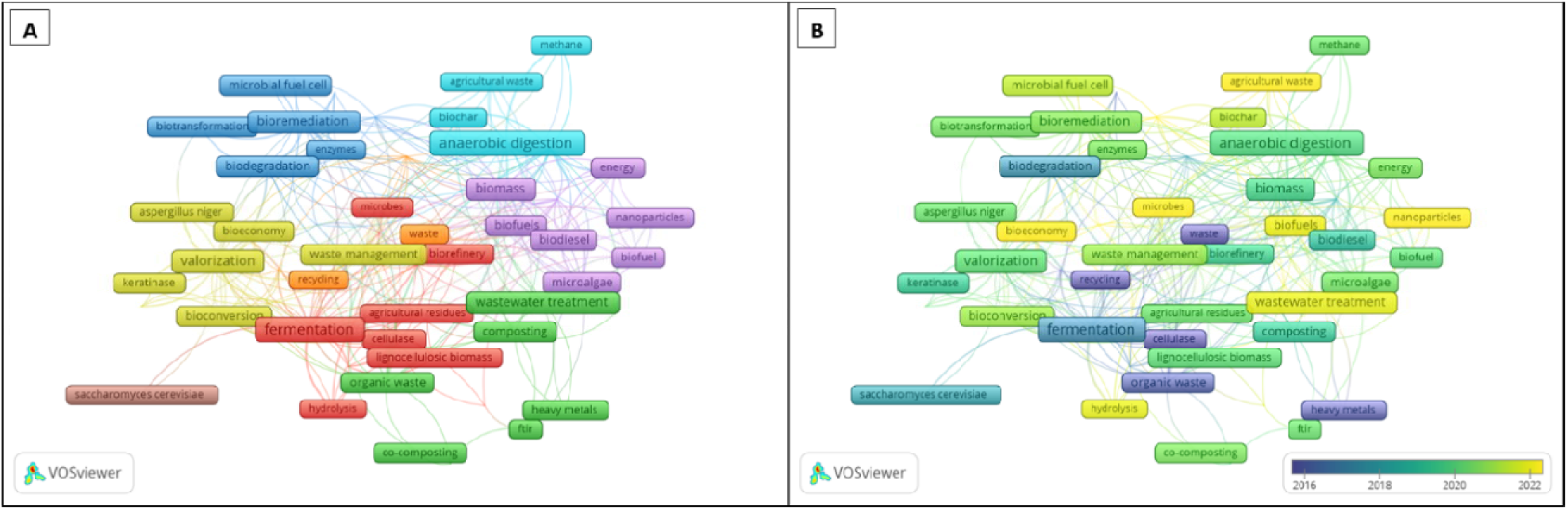
Author Keywords Co-occurrence Network (A) and overlay by average yearly publication (B) from microbial-driven waste Biomass Valorization Research in Africa.

#### 3.9.3 Co-occurrence—index keywords

Out of 5058 index keywords listed by Scopus on this theme, 525 keywords with a total link strength of 84151 were selected based on 5 minimum co-occurrence. Figure 18 depicts a network of index keywords co-occurrence (A) and overlay visualization of index keywords co-occurrence by average yearly publication (B) in microbial-driven waste biomass valorization research in Africa. There are 5 clusters highlighted in the index keywords co-occurrence.Cluster 1 (red frames) had 131 keywords out of the 525 indexed keywords selected. The green frames represent the second cluster with 122 index keywords. Cluster 3, with 112 items, Cluster 4 depicted by the yellow color frames consists of 111 index keywords, while cluster 5 represented by purple frames contains 49 index keywords.

**Figure 18:**
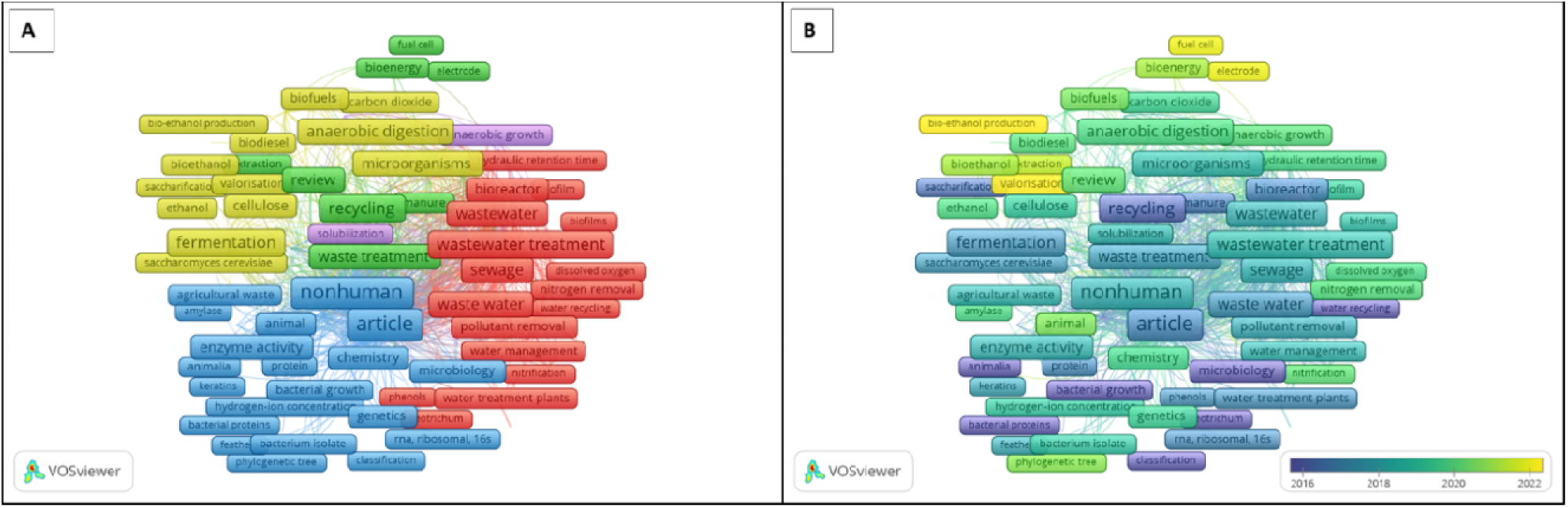
Index keywords Co-occurrence Network (A) and overlay by average yearly publication (B) from microbial-driven waste Biomass Valorization Research in Africa.

## 4.0 Discussion

The high annual growth rate per document of 11.13% in microbial-driven waste biomass valorization research in Africa in comparison to the 4.7% average annual growth rate for scientific publications (Larsen and von, 2010), as well as the consistent increase in publication around this subject in the last 2 decades are indications of the growing interest in waste valorization via the activity of microorganisms. Although the number of publications on microbial-driven waste biomass valorization with regards to Africa were observed to be redundant and irregular within certain periods, there was an observable moderate increase in the number of publications across the three periods. The high percentage increase in the average yearly publication in the rapid growth period confirms that there has been a growing interest in this area of research in Africa; probably due to the inclusion of sustainable cities and communities (SDG 11), as well as responsible production and consumption (SDG 12) in the SDG’s in 2015 (Pérez-Escamilla, 2017). The average yearly citations of documents on microbial-driven waste biomass valorization in Africa fluctuated across the classified periods, with the highest recorded in 2005 unlike what was expected based on the average yearly publications. The reason for this high citation index may be attributed to the quality of the work from Hamdi and Bouallagui research groups’ judging by the fact that there were only two (2) publications in that year which ended the era. Thus, there may have been an increased use of this work by Hamdi and Bouallagui on “Bioreactor performance in anaerobic digestion of fruit and vegetable wastes” in the research done after this period. The total citations till date is above 12,480, indicating an expanding accumulation of knowledge about microbial-driven waste biomass valorization in Africa. Notwithstanding that Nigeria is ahead in terms of the number of publication on microbial-driven waste valorization research in Africa with a total publication of 369, South Africa is far ahead of Nigeria in terms of the rate of document citations; highlighting the impact and depth of the work, as well as the ongoing search for the best waste management techniques in South Africa according to Karani and Jewasikiewitz (2006). Godfrey and Oelofse (2017) reported that South Africa has effectively developed a recycling industry, significantly supported by the diligent efforts of an engaged informal trash sector, over the past three decades. South Africa’s impact in waste valorization by microorganism in Africa is also visibly seen in her collaboration with other countries, with up to 20% inter-country collaborations and 80% intra-country collaborations. Furthermore, 4 of the top 5 most globally referenced documents were from South Africa, with most (40%) of the top 20 authors’ resident in South Africa. Despite this achievement, countries like China, Italy, Malaysia, Canada, Hong Kong, United Kingdom, Germany and Finland were dominant in intercountry collaboration with 100% MCP. Notably, more than 70% of the countries showed more tendencies for inter-country collaborations on microbial-driven waste biomass valorization in Africa. In terms of journal ranking for researchers in the area of microbial-driven waste valorization in Africa, Bioresource Technology journal ranked first in the choice of journals for publication on waste valorization by microorganisms with more than 17% published documents. This high publication index by may be attributed to the scope of the journal, which provides international coverage on the advance and distribution of information in all the related areas of biomass, biological waste treatment, bioenergy, biotransformations, and bioresource systems analysis, as well as technology involved with conversion or generation.

Author’s significance over time, determined by their production and influence within a specific topic area (Forliano et al., 2021, Fadiji et al., 2023) was assessed. The top 3 leading authors (Hamdi M, Hafidi M, and Bux F) in microbial-driven waste valorization research in Africa according to number of publications within the period considered were from Tunisia and South Africa, with a total of 34 publications. For 23 years, Hamdi M publication productivity and output was approximately 1.69 total citations starting from the year of 2000 which had only one publication. However, the 2005 publication on the performance of bioreactor in anaerobic digestion of waste from fruits and vegetables (FVW), enhanced his productivity and output view due to the sharp increase in the total yearly citations. The leadership of Hamdi M was also confirmed based on four criteria used to ascertain the local impact of the authors in waste biomass valorization research in Africa for the top 20 authors. The criteria include how many times the author was cited (TC), the h-index (metric measurement at the author level that quantifies the productivity and citation effect of a scientist’s or scholar’s writings), the g-index (An author level metric that measures the unique largest number of articles published by an author which has been cited for g times such that the top g articles altogether received at least a square of the g citations when a set of articles published are ranked in the decreasing order of citations received) (Egghe, 2006), and the m-index (a time-adjusted evaluation of a researcher’s output). The h-index quantitatively measures an author’s publication volume relative to the citations received. This statistic evaluates the comprehensive influence of an author’s academic contributions and effectiveness. High-cited papers are given more weight by the g-index; as it gauges the distribution of citations received by a specific researcher’s publications; while the m-index, somewhat measures the yearly h-index from the first publication as it is gotten by dividing the h-index by the active years of a researcher (Hirsch, 2007; Forliano et al., 2021; Fadiji et al., 2023). Hamdi M had an h-index, g-index, and total citation (TC), with 12, 15, and 1003 h-index, g-index and total citation (TC) respectively, notwithstanding the lower m-index (0.48) after 25 years (supplementary table S1). The author with the highest m-index (1.667) was Armegam N with a total citation of 276 after 3 years. Hafidi M on the other hand with output and publication productivity spanning over a 19 years; had the most visible outlook in year 2019 with 3 publications and 12 yearly citations, followed by Bux F, with the highest number of articles recorded in 2013 and 2021 with 10.0 and 10.31 yearly citations respectively. Detailed report from the work by Hamdi M in 2005 concluded that continuous two-phase systems were proposed to be the more highly efficient technologies for anaerobic digestion of FVW. One of the major limitations of anaerobic digestion of fruit and vegetable wastes as reported in their work is a rapid wastes acidification which reduces the pH in the reactor, which leads to larger volatile fatty acids production (VFA), stress and inhibition of methanogenic bacteria (Bouallagui et al., 2005). Whereas, the three works by Hafidi M in 2019 with citation per year of 6.71, 1.29, and 4.00 where on “the fate of antibiotics found in a wastewater treatment plant’s primary sludge when co-composted with palm waste; the evaluation of compost-derived humic acids structure through TMAH-Thermochemolysis on lignocellulose waste; and the industrial-scaling of green waste composting using a GORE(R) cover membrane focusing on the physico-chemical characteristics and spectroscopic evaluation” (Khadra et al., 2019; El Ouaqoudi et al., 2019; Al-Alawi et al., 2019). There are serious risks to the environment and public health from the emergence and spread of antibiotic resistance in wastewater. In natural environments, bacteria that carry several antibiotic resistance genes (ARGs), especially those linked to horizontal gene transfer (HGT), can act as long-lasting reservoirs and carriers of antimicrobial resistance (Li et al., 2025).

Recent work by Hafidi in 2024 explored on a long term basis the impact of repetitive application of urban sewage sludge on a sandy soil properties, focusing on the spread of bacteria resistant to antibiotics, metal, and their corresponding resistance genes in soil (Mokni-Tlili et al., 2024). The outcomes of their work showed a dose-dependent variation of different soil parameters like the heavy metal content, total heterotrophic bacteria (THB) etc. Furthermore, the high citation of the work by Hamdi and colleagues in 2005 with 557 citations confirmed their contributions in microbial-driven waste biomass valorization in Africa. Although this could be judged differently based on the age of the publication. In their work, they observed that a continuous two-phase system might be the better technological approach for FVW anaerobic digestion; as about 70– 95% of organic matter was converted to methane at a 1–6.8 g versatile solids per volume of organic loading rate (OLR) (VS)/l in a day. However, Amobonye et al. (2020) highlighted in their review the multifaceted mechanisms and roles microorganisms like bacteria, actinomycetes, fungi, algae and their enzymes play in the degradation of synthetic plastics. Notably, another work by Amobonye et al. (2021), on “Plastic biodegradation: frontline microbes and their enzyme” with a total citation of 414, and 3rd in the rank of top 11 most cited documents based on the total citation per document, had the highest yearly mean citations of 103.5 after 4 years. The significant interest in plastic bioremediation could be associated with the quest for the elimination of plastics and microplastics, which have presented substantial environmental and health hazards in recent years. Yang et al. (2020) emphasized that over the previous 65 years, cumulative plastic trash has accumulated to around 6300 million metric tons (Mt). Plastics are broken down by microorganisms into microscopic particles known as microplastics (MPs). According to reports, MPs carry organic pollutants like pesticides, antibiotics, and polybrominated diphenyl ethers, which can result in intricate interactions (Yang et al., 2020). Following the publication by Bouallagui and colleagues as second most cited documents is the work by Ayilara et al. (2020) with a 490 total citations and a yearly mean citations of 98. Ayilara et al. (2020) worked on the challenges and potentials of using composting in waste management, where they highlighted that the advantage in the use of compost is the ability of compost to improve soil structure and nutrient availability, thus increasing the nutrient already available. The work by Olaniran et al. (2011) on “Chlorophenols and other related derivatives of environmental concern: Properties, distribution and microbial degradation processes” occupied the 4th spot based on the 400 total accumulated citations over a 14 years period accruing to a yearly mean citation of 28.57. the remaining 6 spots were occupied by Jimoh et al. (2019), Beyene et al. (2018), Annab et al. (2007), Kizito et al. (2017), Mnif and Ghribi (2016), Wu et al. (2016), and Nsenga et al. (2019) with a total citation and mean yearly citations of 243 and 40.5, 213 and 30.43, 211 and 11.72, 191 and 23.88, 178 and 19.78, 166 and 18.44, and 105 and 17.5 respectively. Of all the 11 most cited documents, 1 (9.09%) was a research article, others 10 (90.01%) were review articles. The dominance of Review articles in microbial driven waste valorization research in Africa highlights the depth of interest and confidence researchers have for Review articles. Miranda and Garcia-Carpintero (2018) reported that Review articles are typically cited more often, at least three times more, than standard research papers. This could also be due to the condensation of information from diverse authors on a particular topic by Review articles. Review articles are studies that critique and integrate current literature by identifying, contesting, and enhancing the foundational elements of a theory through the analysis of one or more bodies of past research (Post et al., 2020; Bahishti, 2021). Reviews can facilitate knowledge advancement, establish guidelines for policy and practice, furnish evidence of application, identify research gaps, and, if executed effectively, possess the potential to inspire new ideas and form the foundation for future research trajectories (Bahishti, 2021). Additionally, Review articles are also cited even when the main work cited by the Review article have been explored and cited, thus giving rise to the continuous increase in Review article citation (Bahishti, 2021).

The trending topics in microbial-driven waste valorization research in Africa crisscrossed many aspects, ranging from the valorization of molasses from sugarcane to the use of phanerochaete species, and *Enterobacter aerogenes* in waste management. *Phanerochaete chrysosporium*, a white-rot fungus, has been utilized extensively to clean waste streams that contain hazardous organic contaminants and heavy metals (Zeng et al., 2012). Huang et al. (2008) reported the use of *Phanerochaetechrysosporium* in the reduction of lead toxicity and the degradation of lead-contaminated lignocellulosic waste. *Phanerochaete chrysosporium* under combinatorial stress creates varied oil profiles after secondary metabolite analysis, according to Whiteford et al. (2021). Furthermore, Reungsang and colleagues in 2013 reported that Enterobacter aerogenes KKU-S1 produces hydrogen and ethanol simultaneously from waste glycerol. According to Pachapur et al. (2016), a co-culture system employing Enterobacter aerogenes NRRL B-407 and Clostridium butyricum NRRL B-41122 was used to valorize crude glycerol (CG) for enhanced hydrogen generation. Ethanol production Optimization in acetic acid and glycerol waste co-substrate by Enterobacter aerogenes was the focus of Sawasdee et al. (2021). From 2007 to 2012, there seemed to be increased interest in the search for beneficial uses of agro/industrial wastes and soil microbes. Enterococcus copious in wastewater was reported by Layton et al. (2010) to be drawn from human feces which contain significant amounts of enterococci ranging from 104 to 106 bacteria per gram of wet weight. Benrebah et al. (2007) highlighted the use of wastewater sludge and agro-industrial waste materials for rhizobial inoculant production in their review. Geotrichum candidum, another microorganism of interest during this period, has been reported to have potential in olive mill wastewater treatment (Asses et al., 2009). Both wastes from the processing of cassava (Manihot esculenta) and textile production were also another topic of discussion within 2007 to 2012. From 2013 till 2019, attention began shifting from alcohol production from waste to anaerobic digestion of waste and enzymatic processes in waste management. Hedge et al. (2018), reported on the characteristics and uses of food processing wastes in sustainable alcohol production. Bacillus pumilus and Lactobacillus plantarum were also among the trending organisms with application in wastewater and agro waste management. Mlaik et al. (2014) mentioned the use of two proteolytic microbes (Bacillus pumilus and Bacillus cereus) in the unhairing treatment of wastewater. The combinatorial effect of the use Lactobacillus plantarum MTD1 and waste molasses as fermentation modifier was evaluated by Zhao et al. (2029) and it stated that this combination increased silage quality, as well as reduced ruminal greenhouse gas emissions of rice straw. L. plantarum has also been documented in recent times to improve the efficiency of sheep manure composting and product quality (Li et al., 2019). The usefulness of the earthworm Elsenia fetida in vermicomposting may have enhanced the highlighting of this earthworm in the trending issue within the period of 2013 to 2019. This nematode was reported by Mane et al. (2024) to have improved waste management via the conversion of organic waste into nutrient-rich biofertilizers when it was applied in the management of municipal solid waste. From 2020 till 2024 when this review was done, the trending issues were narrowly around waste valorization as different concepts were seen speaking to these issues. The trending topics ranged from recycling, wastewater treatment, anaerobic digestion, enzymatic hydrolysis, valorization, plastic recycling, purification etc. Furthermore, the thematic keyword trends and map diagram in Figure 14 highlights an analysis of the trend of keyword plus in microbial-driven waste biomass valorization research in Africa. Thematic trend maps are an essential analysis in bibliometric reviews because they tend to highlight the historical aspects of research move from the non to the unknown, marking the path to future perspectives (Chen et al., 2019; Chansanam and Li, 2022; Fadiji et al., 2023). The analysis was conducted using the “Walktrap” clustering algorithm, with 1,000 words, a minimum cluster frequency (per thousand documents), and five labels. The thematic trend map consists of four quadrants depicting the four levels and degree of relevance of these keywords in the past, currently and the future. The lower left quadrant (emerging and declining themes) encompasses subjects characterized by low density and focus, indicative of themes that have recently emerged or vanished. Two clusters are shown in this quadrant based on the color. The light green cluster consisting of enzyme activity, temperature, etc. tends to reveal the declining themes, while the light blue cluster consisting of biomass, anaerobic digestion, etc tends to show emerging themes as corroborated by trending topics and thematic evolution in figure 12 and 14 respectively. The region in the upper left quadrant (Niche themes) denotes a specific topic mostly excluded from the primary emphasis which could be said to be a niche in research on microbial-driven waste biomass valorization. The regions in the upper right quadrant (motor themes) signify well-established and resilient subjects within the domain of research on microbial-driven waste biomass valorization. This quadrant contains two clusters signified with pink and purple colors. The pink color cluster reveals highly relevant themes with slow degree of development; whereas the purple cluster shows themes that are of seldom relevance and high developmental degree. The regions in the lower right quadrant denote the most prevalent fundamental and transverse subjects, however internal development is not as enhanced as that of the upper right quadrant. The themes in the lower left and right quadrants may serve as a source of inspiration and potential reference for more research in microbial-driven waste biomass valorization.

When considering the theme evolution using keyword plus (keywords based on datasets), keywords like cellulase, biodegradation, anaerobic digestion, waste and waste water management were quite popular in the first period (1984–2005). These keywords subsequently evolved into the next period (2006–2016) as biomass, wastewater, and animal. The keywords in the second period metamorphosed into nonhuman, wastewater treatment and anaerobic digestion in the last period (2017-2024). The term article in the first and second period may be speaking to the primary source of data for bibliometric analysis, as articles are the primary unit of academic publishing. The article in the second period originated primarily from cellulose and biodegradation, aside from the original conceptualization in the first generation. The keywords biomass and wastewater in the second period were observed to have stemmed mainly from keywords like article, biodegradation, waste management, anaerobic digestion, and waste water management in the first period. Some aspects of all the keywords in the second period had a merger into nonhuman in the last period. When discussing waste valorization, the term “nonhuman” is often used to refer to waste generated by nonhuman sources, such as agricultural residues, industrial by-products, or animal waste, which can be transformed into valuable products like fertilizers, biofuels, or bioplastics using a range of valorization techniques. For example, crop residues and food processing waste are nonhuman sources with significant potential for sustainability in material development. This trend demonstrates that waste and waste water management, and anaerobic digestion are the core topics in the research on microbial-driven waste biomass valorization in Africa judging by the appearance of these keywords in the first and last periods of consideration in this study.

Keywords (most re-occurring words that characterize a literature) were also used to understand the type of information retrieved from diverse publications on the subject matter. According to Rejeb et al. (2021) and Zhong et al. (2021), keyword co-occurrence is most established in at least two publications in a period. However, a minimum of 5 keyword occurrence was adopted, which resulted in only 570 keywords meeting the threshold out of the 5833 keywords listed. The keywords are arranged in 7 clusters signified by the frames with different colorations and sizes. The clusters and colorations are sometimes determined either by closeness of their usage, or by the average publication per year and/or average citation per year. The frames are connected to each other depending on the strength of their cooperative use, and this is known as the link strength (the frequency of the keyword’s co-occurrence). The total link strength is the summation of all the keywords link strengths over all the other keywords (Guo et al., 2019; Martynov et al., 2020). The size and the distance separating the frames corresponds to the corresponding frequency of co-occurrence of the keyword in publications and the density which impacts the visualization according to Rejeb et al. (2021). Figure 16a showed all author keywords with Network visualization. The cluster 1 represented by the red framed keywords has 135 keywords clustered in it with fermentation, biomass, and anaerobic digestion the three most highly used keywords in this cluster having an average yearly citations (ayc) of 26.41, 22.88, 31.12 (fig. 16b) and occurrence frequency (fc) of 83, 72, 69 respectively. Cluster 2, represented by green, contained 102 keywords, with waste treatment, microbial community, and composting as the three highest occurring keywords with fc and ayc of 50 and 31.18, 38 and 34.82, and 38 and 41.16 respectively. Clusters 3 and 4, represented by blue and yellow, have 100 and 99 keywords clustered in them, with recycling, bioremediation and review having the three (72, 47, and 47) highest fc with ayc of 23.47, 47.98, and 65.51 respectively in cluster 3. Cluster 4 has nonhuman, controlled study, and pH as the three most occurring keywords with 152, 65 and 61 fc respectively. Finally, cluster 5, 6 and 7, represented by purple, turquoise blue and orange color respectively contains 98, 28 and 12 keywords with the most occurring keywords in these cluster being wastewater treatment, microalgae, and anaerobiosis having fc of 72, 23 and 18 respectively. In clusters 5 and 6 keywords of the same meaning and origin but written differently were classified differently. For instance in cluster 5 “wastewater” and “waste water” classified differently (62 and 52 fc respectively) would have had the highest fc had the two keywords been recognized as one word. “Microalgae”, “microalga” and “micro-algae” are three keywords recognized differently which had a total fc and ayc of 61 and 68.75 respectively. Keyword co-occurrence network aids in the identification of the main content and knowledge framework of a publication (Fadiji et al., 2023). Using authors provided keywords, a total of eight (8) clusters having a total link strength of 363 and distinct colors were observed in the co-occurrence network (Figure 17a). Cluster 1 (red framed keyword), 2 (green framed keyword) and 3 (blue framed keyword) each consist of 10 keywords. Cluster 1 and 2 has words like fermentation and composting earlier reported in the all author keywords also having the highest occurrence of 33 and 19 respectively. Thus showing the significance of such words and systems in microbial-driven waste biomass valorization. In cluster 3, bioremediation with 19 fc proved to be the highest co-occurring keywords in this cluster, highlighting the interest of many authors in the use of biomaterials for the cleaning and valorization of waste. The yellow color framed keywords represents cluster 4 (9 keywords) and has valorization with 24 occurrences and 28 link strengths as the highest occurring keyword. Although, with the combination of valorization and valorisation, the total link strength was 34 while the fc was 30. Other keywords in this cluster are bioconversion, bioeconomy, enzyme, keratinase, waste management, etc. Cluster 5 represented in purple contains 8 keywords which includes wastewater, biomass, biofuel(s), biodiesel, energy, nanoparticles, and microalgae. The highest keywords in this cluster were biomass and wastewater, which may be attributed to the topic of “waste biomass valorization and clean environment associated with number 6, 11 and 12 of the united nation sustainable development goal.” Clusters 6 (5 keywords), 7 (4 keywords) and 8 (1 keyword), represented by torques blue, orange and brown colors, have anaerobic digestion (32 fc and link strength) and microorganisms (12 fc and 18 link strength) as the highest occurring words in cluster 6 and 7 respectively. Other keywords contained in clusters 5 and 6 are agricultural waste, biogas, methane and biochar, and biotechnology, recycling and waste respectively. Saccharomyces cerevisiae is the only keyword contained in cluster 8. More than 50% (29) of the categorized keywords based on the 5 co-occurrence acceptability were from 2020 to 2022. Aside 2016 and below when 5.26% co-occurrence was recorded, all the other years within which the re-occurring keywords were acceptable had more than 10% co-occurrence (figure 17b; supplementary table 2). On the other hand, using indexed keywords from Scopus database on the subject of microbial-driven waste biomass valorization research in Africa, there was only Five (5) clusters with a total 525 indexed keywords and a total link strength of 84151 selected based on 5 minimum co-occurrence standard. Index keywords are selected keywords from Scopus database which are standardized based on the vocabularies of that thematic focus licensed or owned by Elsevier and derived from thesauri (Fadiji et al., 2023). This set of keywords is slightly different from all keywords and author keywords as it considers alternative spellings, synonyms and plurals according to Golub et al. (2020). Cluster 1 (red frames) with 131 keywords out of the 525 indexed keywords selected, contains Keywords like sewage, waste water, wastewater, waste water management, wastewater treatment, etc. with the latter having the highest (75) frequency of occurrence and link strength (1983). The sum of the link strength and frequency from the aforementioned keywords which were the top 5 keywords based on the frequency of occurrence were 8070 and 299 respectively. Contributing up to 9.6% of the total link strength of the 525 index keywords. The green frames represent the second cluster with 122 index keywords. The 3 top ranking keywords in this cluster based on the frequency of occurrence are recycling (72), waste treatment (50) and review (46); contributing up to 4.5% of the total link strength. Cluster 3, with 112 items in blue, has article and nonhuman as keywords with the highest (152) occurrence. Other keywords in cluster 3 are controlled study, bacteria, bacterium, biodegradation, agricultural waste, agro-industrial waste etc. Cluster 4 depicted by the yellow color frames consists of 111 index keywords with words like fermentation, biomass, anaerobic digestion being the top 3 most occurring keywords with 66, 66, and 55 occurrences respectively. The cluster 5 represented by purple frames is the last cluster in the network co-occurrence with 49 index keywords including words like microbial community, anaerobic growth, anaerobiosis, methane, etc. The keywords with frequent occurrences beyond the 2022 publication year from figure 18B were words like energy, bioenergy, pollutants, electron transport, bio ethanol, bio-ethanol productions, etc. This could be speaking to the current and future perspectives in the field of microbial-driven waste biomass valorization research.

### 4.1 Future perspective: Harnessing the power of anaerobic digestion and biofilms in waste management and plastic recovery

Waste prevention, recycling, reuse, and recovery are important waste management strategies that ease the burden on landfills, conserve natural resources, and save energy. Implementing a combination of these strategies can improve waste management sustainably (Wan et al., 2019). Anaerobic digestion is a sustainable process for processing residues and wastes in the agro-food industry, offering the potential to treat biodegradable wastes or produce valuable products. It can help in managing solid organic waste sustainably to avoid resource depletion and reduce environmental burdens. Anaerobic digestion has the potential to convert raw solid organic wastes into useful products like biogas and energy-rich compounds, which can contribute to meeting the increasing global energy demands in the future (Ibrahim et al., 2016). Anaerobic digestion (AD) significantly impacts waste management by offering a sustainable solution for treating various organic waste streams. It involves the breakdown of organic matter in the absence of oxygen by a diverse community of microorganisms, resulting in the production of biogas (primarily methane and carbon dioxide) and digestate (a nutrient-rich byproduct). This process offers several advantages such as (i) Waste Reduction and Volume Minimization, in which the volume of organic waste sent to landfills is reduced, thus decreasing greenhouse gas emissions associated with landfilling and conserving landfill space. (ii) Renewable Energy Generation: Biogas produced can be used to generate electricity, heat, or biofuels, contributing to renewable energy production and reducing reliance on fossil fuels. (iii) Resource Recovery: Digestate, a byproduct of AD, is a valuable fertilizer rich in nutrients like nitrogen, phosphorus, and potassium, which can be used in agriculture, reducing the need for synthetic fertilizers. (iv) Greenhouse Gas Emission Reduction: By diverting organic waste from landfills, AD significantly reduces methane emissions, a potent greenhouse gas. Methane produced during AD can be captured and used, preventing its release into the atmosphere. Future research in waste valorization through microbial processes be focused on improving AD efficiency via the optimization of process parameters such as temperature, pH, retention time, and inoculum selection to enhance biogas yield and reduce process inhibition; as well as developing advanced pretreatment techniques to enhance the digestibility of recalcitrant organic materials. Co-digestion of different waste streams is another possible leap in AD improvement strategies, thus improving digestate quality and biogas yield. Other perspectives are towards developing novel bioprocesses for waste valorization by exploring microbial communities and metabolic pathways for the production of high-value bioproducts (e.g., bioplastics, biofuels) from waste.

Microbes also play a crucial role in plastic recovery by breaking down polymer bonds to produce smaller components that can be further processed. Various solutions such as waste sorting, value-added recycling, energy recovery, product bans, and biodegradable plastic production have been proposed to address plastic waste challenges (Tamoor et al., 2021).

Bacterial biofilms have emerged as a promising strategy for the valorization of food waste, converting diverse food substrates into valuable products like biofuels, enzymes, and biopolymers (Das et al., 2024, Sarkar et al., 2025). Functional biofilms have been evidenced in literature for several applications, including catalysis, electrical conduction, bioremediation, and medical therapy. The mechanical properties of biofilms can be modified through genetic editing, metal ion curing, and synthetic gene circuits, among other methods (Li et al., 2022).

## 5.0 Conclusion

Microorganisms with their complex biochemical functionaries play a significant role in waste valorization. Research in the application of microorganisms in waste conversion has received tremendous attention over many decades. The use of bibliographic data from Scopus database to study the extent of research in this area in Africa was significant as information regarding the top ranked authors and institutions, highly cited documents and trending topics and themes, as well as future perspectives in this field were revealed. Hamdi M, Hafidi M, and Bux F were the leading top 3 authors as Nigeria ranked above other countries from 1984-2024 based on the number of articles published. Although, in terms of number of citations from a single country publication (SCP) South Africa dominated with 3233 and 75% SCP. The top 3 most important journal sources were Bioresources Technology, Journal of Environmental Management, and Science of the Total Environment cumulatively boasting. Themes like wastewater treatment, composting, recycling, and microorganisms evolved to biodegradation, biomass, microbial community, anaerobic digestion etc. The findings suggest that emerging themes are still revolving around biomass and anaerobic digestion, as more research is still ongoing in wastewater treatment and management, recycling and fermentations in waste valorization. This knowledge contributes to future research designing and gap identification in the area of waste valorization and plastic recovery in Africa.

---

**Figure S1:**
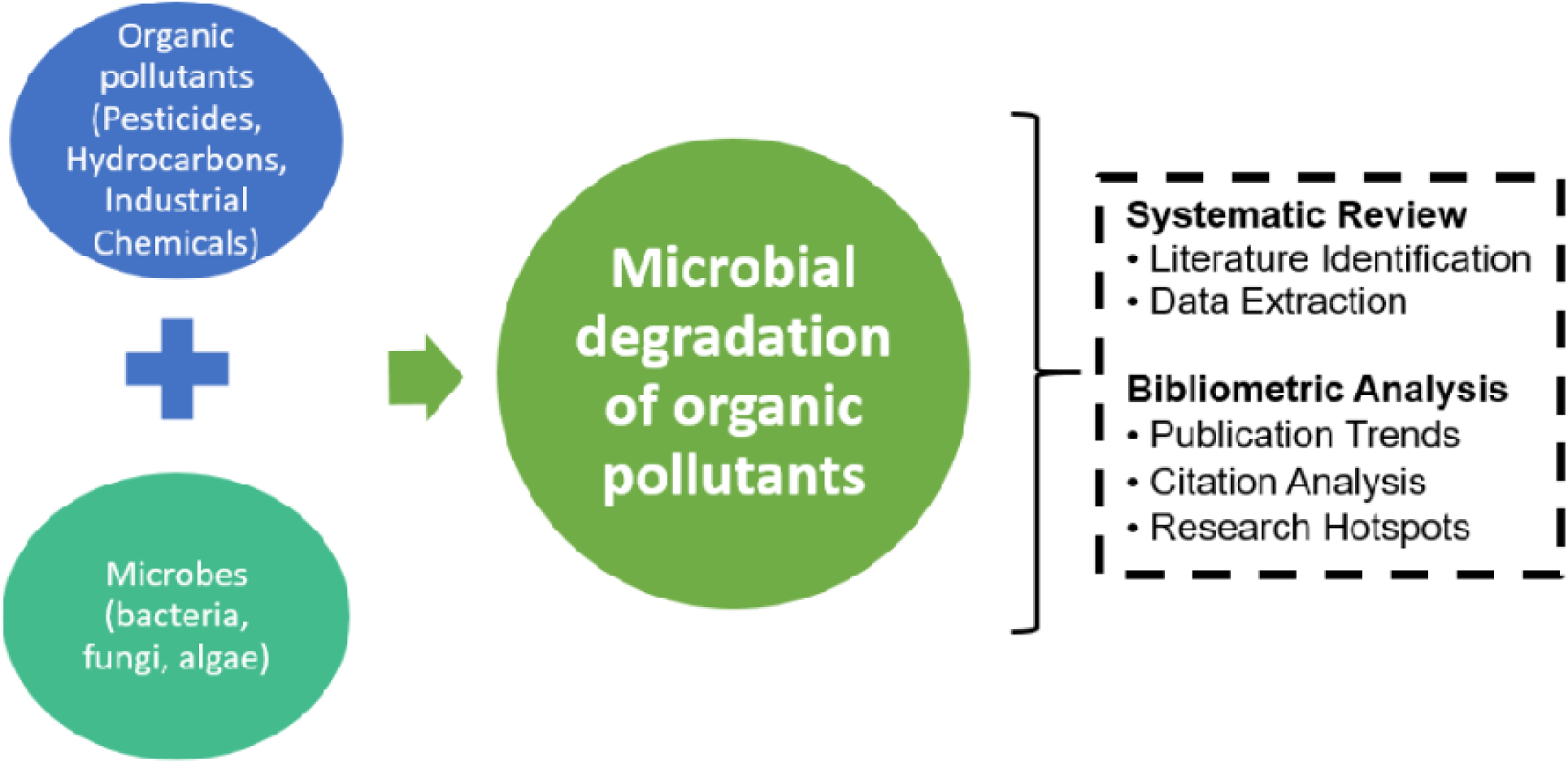
Schematic representation of the major concepts in this study.

**Supplementary Table S1:** Authors Productivity and impact in Research on microbial-driven waste Biomass Valorization in Africa.

| Author | h-index | g-index | m-index | TC | NP | PY-start |
| --- | --- | --- | --- | --- | --- | --- |
| **Hamdi M (Tunisia)** | 12 | 15 | 0.48 | 1003 | 15 | 2000 |
| **Bux F (South Africa)** | 7 | 9 | 0.438 | 279 | 9 | 2009 |
| **Hafidi M (Tunisia)** | 7 | 10 | 0.368 | 173 | 10 | 2006 |
| **Nwodo UU (South Africa)** | 7 | 8 | 1.167 | 175 | 8 | 2019 |
| **Nnolim NE (South Africa)** | 6 | 7 | 1.2 | 157 | 7 | 2020 |
| **Bouallagui H (Tunisia)** | 5 | 5 | 0.25 | 775 | 5 | 2005 |
| **Karmegam N (India)** | 5 | 5 | 1.667 | 276 | 5 | 2022 |
| **Okoh AI (South Africa)** | 5 | 5 | 0.833 | 140 | 5 | 2019 |
| **Sillanpää M (South Africa)** | 5 | 5 | 1.25 | 90 | 5 | 2021 |
| **Van Zyl WH (South Africa)** | 5 | 7 | 0.455 | 108 | 7 | 2014 |
| **Dahunsi SO (Nigeria)** | 4 | 5 | 0.571 | 120 | 5 | 2018 |
| **Ekama GA (South Africa)** | 4 | 4 | 0.444 | 297 | 4 | 2016 |
| **Kasmi M (Tunisia)** | 4 | 5 | 0.5 | 73 | 5 | 2017 |
| **Lateef A (Nigeria)** | 4 | 4 | 0.571 | 119 | 4 | 2018 |
| **Li X (China)** | 4 | 5 | 0.667 | 109 | 5 | 2019 |
| **Lyamlouli K (Morocco)** | 4 | 6 | 1 | 63 | 6 | 2021 |
| **Nasri M (Tunisia)** | 4 | 4 | 0.286 | 159 | 4 | 2011 |
| **Ouhdouch Y (Morocco)** | 4 | 5 | 1 | 48 | 5 | 2021 |
| **Sayadi S (Tunisia)** | 4 | 4 | 0.211 | 118 | 4 | 2006 |
| **Sithole B (South Africa)** | 4 | 4 | 0.571 | 88 | 4 | 2018 |

**Supplementary Table 2:** percentage occurrence of author keyword by average yearly publication.

| Publication year range | Number of occurrence | Occurrence (%) |
| --- | --- | --- |
| above 2022 | 6 | 10.53 |
| 2020-2022 | 29 | 50.88 |
| 2018-2020 | 9 | 15.79 |
| 2016-2018 | 10 | 17.54 |
| below 2016 | 3 | 5.26 |

## References

Abraham, P., Chinedu, E. O., Banaru, A. A., & Dauda, W. P. (2023). Biosurfactants: possible roles in environmental management-a review. FUDMA JOURNAL OF SCIENCES, 7(4), 236–245.

A.K. Ziraba, T.N. Haregu, B. Mberu, A review and framework for understanding the potential impact of poor solid waste management on health in developing countries, Arch. Publ. Health 74 (55) (2016) 1–11, 10.1186/s13690-016-0166-4. https://archpublichealth.biomedcentral.com/track/pdf/10.1186/s13690-016-0166-4.pdf.

African Clean Cities Platform (ACCP), Basics of Municipal Solid Waste in Africa, 2019. https://unhabitat.org/african-clean-cities-publications.

Afrifa, S.; Zhang, T.; Appiahene, P.; Varadarajan, V. Mathematical and Machine Learning Models for Groundwater Level Changes: A Systematic Review and Bibliographic Analysis. Future Internet 2022, 14, 259. 10.3390/fi14090259

Al Ryalat, S.A.; Malkawi, L.W.; Momani, S.M. Comparing bibliometric analysis using PubMed, Scopus, and Web of Science databases. JoVE (J. Vis. Exp.) 2019, 152, e58494.

Al-Alawi, M., Szegi, T., El Fels, L., Hafidi, M., Simon, B., & Gulyas, M. (2019). Green waste composting under GORE(R) cover membrane at industrial scale: physico-chemical properties and spectroscopic assessment. International Journal of Recycling of Organic Waste in Agriculture. doi:10.1007/s40093-019-00311

Alfadil, M.O.; Kassem, M.A.; Ali, K.N.; Alaghbari, W. Construction Industry from Perspective of Force Majeure and Environmental Risk Compared to the COVID-19 Outbreak: A Systematic Literature Review. Sustainability 2022, 14, 1135.

Amobonye, A., Bhagwat, P., Singh, S., & Pillai, S. (2020). Plastic biodegradation: frontline microbes and their enzymes. Science of The Total Environment, 143536. doi:10.1016/j.scitotenv.2020.1435

Annabi, M., Houot, S., Francou, C., Poitrenaud, M., & Bissonnais, Y. L. (2007). Soil Aggregate Stability Improvement with Urban Composts of Different Maturities. Soil Science Society of America Journal, 71(2), 413. doi:10.2136/sssaj2006.0161

Asses, N., Ayed, L., Bouallagui, H., Benrejeb, I., Gargouri, M., & Hamdi, M. (2009). Use of Geotrichum candidum for olive mill wastewater treatment in submerged and static culture. Bioresource Technology, 100(7), 2182– 2188. doi:10.1016/j.biortech.2008.10.04

Ayilara, M., Olanrewaju, O., Babalola, O., & Odeyemi, O. (2020). Waste Management through Composting: Challenges and Potentials. Sustainability, 12(11), 4456. doi:10.3390/su12114456

Bahishti, A. A. (2021). The importance of review articles & its prospects in scholarly literature. Extensive Reviews, 1(1), 1–6. 10.21467/exr.1.1.4293

Benrebah, F., Prevost, D., Yezza, A., & Tyagi, R. (2007). Agro-industrial waste materials and wastewater sludge for rhizobial inoculant production: A review. Bioresource Technology, 98(18), 3535–3546. doi:10.1016/j.biortech.2006.11.06

Beyene, H. D., Werkneh, A. A., & Ambaye, T. G. (2018). Current updates on waste to energy (WtE) technologies: a review. Renewable Energy Focus, 24, 1–11. doi:10.1016/j.ref.2017.11.001

Bouallagui, H., Touhami, Y., Ben Cheikh, R., & Hamdi, M. (2005). Bioreactor performance in anaerobic digestion of fruit and vegetable wastes. Process Biochemistry, 40(3-4), 989–995. doi:10.1016/j.procbio.2004.03.007

Chansanam, W., and Li, C. (2022). Scientometrics of poverty research for sustainability development: trend analysis of the 1964–2022 data through Scopus. Sustainability 14:5339. doi: 10.3390/su14095339

Chen, X., Lun, Y., Yan, J., Hao, T., and Weng, H. (2019). Discovering thematic change and evolution of utilizing social media for healthcare research. BMC Med. Inform. Decis. Mak. 19, 39–53. doi: 10.1186/s12911-019-0757-4

Dauda, W. P., Glen, E., Abraham, P., Adetunji, C. O., Morumda, D., Abraham, S. E., … & Ogwuche, I. O. (2023). ELUCIDATING THE FUNCTIONAL ANNOTATION AND EVOLUTIONARY RELATIONSHIPS OF CYTOCHROME P450 GENES IN XYLARIA SP. FL1777 USING IN-SILICO APPROACHES. FUDMA JOURNAL OF SCIENCES, 7(4), 246–264.

Dladla, I., Machete, F., & Shale, K. (2016). A review of factors associated with indiscriminate dumping of waste in eleven African countries. African Journal of Science, Technology, Innovation and Development, 8(5–6), 475–481. 10.1080/20421338.2016.1224613

Drissi, N.; Ouhbi, S.; Marques, G.; Díez, I.d.; Ghogho, M.; Idrissi, M.A.J. A Systematic Literature Review on e-Mental Health Solutions to Assist Health Care Workers during COVID-19. Telemed. e-Health 2021, 27, 594– 602.

Egghe, Leo (2006) Theory and practise of the g-index, Scientometrics, 69(1): 131–152. doi:10.1007/s11192-006-0144-7

El Ouaqoudi, F. Z., Meddich, A., Lemée, L., Amblès, A., & Hafidi, M. (2019). Assessment of Compost-Derived Humic Acids Structure from Ligno-Cellulose Waste by TMAH-Thermochemolysis. Waste and Biomass Valorization. doi:10.1007/s12649-018-0268

Fadiji T, Bokaba T, Fawole OA and Twinomurinzi H (2023) Artificial intelligence in postharvest agriculture: mapping a research agenda. Front. Sustain. Food Syst. 7:1226583. doi: 10.3389/fsufs.2023.1226583

Forliano, C., De Bernardi, P., and Yahiaoui, D. (2021). Entrepreneurial universities: a bibliometric analysis within the business and management domains. Technol. Forecast. Soc. Chang. 165:120522. doi: 10.1016/j.techfore.2020.120522

García-Sánchez, P., Mora, A. M., Castillo, P. A., & Pérez, I. J. (2019). A bibliometric study of the research area of videogames using Dimensions.ai database. Procedia Computer Science, 162, 737–744. 10.1016/j.procs.2019.12.045

Godfrey, L., Tawfic Ahmed, M., Giday Gebremedhin, K., H.Y. Katima, J., Oelofse, S., Osibanjo, O., … H. Yonli, A. (2020). Solid Waste Management in Africa: Governance Failure or Development Opportunity? IntechOpen. doi: 10.5772/intechopen.86974

Golub, K., Tyrkkö, J., Hansson, J., and Ahlström, I. (2020). Subject indexing in humanities: a comparison between a local university repository and an international bibliographic service. J. Doc. 76, 1193–1214. doi: 10.1108/JD-12-2019-0231

Guo, Y. M., Huang, Z. L., Guo, J., Li, H., Guo, X. R., and Nkeli, M. J. (2019). Bibliometric analysis on smart cities research. Sustainability 11:3606. doi: 10.3390/su11133606

Hegde, S., Lodge, J. S., & Trabold, T. A. (2018). Characteristics of food processing wastes and their use in sustainable alcohol production. Renewable and Sustainable Energy Reviews, 81, 510–523. doi:10.1016/j.rser.2017.07.012

Hernández-Torrano, D.; Ibrayeva, L. Creativity and education: A bibliometric mapping of the research literature (1975–2019). Think. Ski. Creat. 2020, 35, 100625.

Hirsch, J. E. (2007). Does the h index have predictive power?. Proceedings of the National Academy of Sciences, 104, 19193–19198.

Huang, D. L., Zeng, G. M., Feng, C. L., Hu, S., Jiang, X. Y., Tang, L., Su, F. F., Zhang, Y., Zeng, W., & Liu, H. L. (2008). Degradation of lead-contaminated lignocellulosic waste by Phanerochaete chrysosporium and the reduction of lead toxicity. Environmental science & technology, 42(13), 4946–4951. 10.1021/es800072c

Jimoh, A. A., & Lin, J. (2019). Biosurfactant: A new frontier for greener technology and environmental sustainability. Ecotoxicology and Environmental Safety, 184, 109607. doi:10.1016/j.ecoenv.2019.109607

Kabongo, J.D. (2013). Waste Valorization. In: Idowu, S.O., Capaldi, N., Zu, L., Gupta, A.D. (eds) Encyclopedia of Corporate Social Responsibility. Springer, Berlin, Heidelberg. 10.1007/978-3-642-28036-8_680

Khadra, A., Ezzariai, A., Merlina, G., Capdeville, M.-J., Budzinski, H., Hamdi, H., … Hafidi, M. (2019). Fate of antibiotics present in a primary sludge of WWTP during their co-composting with palm wastes. Waste Management, 84, 13–19. doi:10.1016/j.wasman.2018.11.009

Kizito, S., Lv, T., Wu, S., Ajmal, Z., Luo, H., & Dong, R. (2017). Treatment of anaerobic digested effluent in biochar-packed vertical flow constructed wetland columns: Role of media and tidal operation. Science of The Total Environment, 592, 197–205. doi:10.1016/j.scitotenv.2017.03.1

Kumari, T., & Raghubanshi, A. S. (2023). Waste management practices in the developing nations: challenges and opportunities. In Elsevier eBooks (pp. 773–797). 10.1016/b978-0-323-90463-6.00017-8

Larsen, P. O., & von Ins, M. (2010). The rate of growth in scientific publication and the decline in coverage provided by Science Citation Index. Scientometrics, 84(3), 575–603. 10.1007/s11192-010-0202-z

Layton BA, Walters SP, Lam LH, Boehm AB. Enterococcus species distribution among human and animal hosts using multiplex PCR. J Appl Microbiol. 2010;109(2):539–574.

Li, W., Liu, Y., Hou, Q., Huang, W., Zheng, H., Gao, X., … Sun, Z. (2019). Lactobacillus plantarum improves the efficiency of sheep manure composting and the quality of the final product. Bioresource Technology, 122456. doi:10.1016/j.biortech.2019.12245

Linnenluecke, M.K.; Marrone, M.; Singh, A.K. Conducting systematic literature reviews and bibliometric analyses. Aust. J. Manag. 2020, 45, 175–194.

Mane, V.B., Kanase, S.S., Sawale, N.S. et al. Application of earthworm Eisenia fetida in waste management in municipal solid waste. Environ Dev Sustain (2024). 10.1007/s10668-024-04974-y

Martynov, I., Klima-Frysch, J., and Schoenberger, J. (2020). A scientometric analysis of neuroblastoma research. BMC Cancer 20, 1–10. doi: 10.1186/s12885-020-06974-3

Miranda, R., & Garcia-Carpintero, E. (2018). Overcitation and overrepresentation of review papers in the most cited papers. Journal of Informetrics, 12(4), 1015–1030. 10.1016/j.joi.2018.08.006

Mishra, D.; Gunasekaran, A.; Papadopoulos, T.; Childe, S.J. Big Data and supply chain management: A review and bibliometric analysis. Ann. Oper. Res. 2018, 270, 313–336.

Mlaik, N., Bouzid, J., Hassan, I. B., Woodward, S., Elbahri, L., & Mechichi, T. (2014). Unhairing wastewater treatment byBacillus pumilusandBacillus cereus. Desalination and Water Treatment, 54(3), 683–689. doi:10.1080/19443994.2014.883330

Mnif, I., & Ghribi, D. (2016). Glycolipid biosurfactants: main properties and potential applications in agriculture and food industry. Journal of the Science of Food and Agriculture, 96(13), 4310–4320. doi:10.1002/jsfa.7759

Mwangomo, E. (2019). Assessment of municipal solid waste management system in Mbeya City Tanzania. International Research Journal of Public and Environmental Health, 6(3). 10.15739/irjpeh.19.006

Nsenga Kumwimba, M., & Meng, F. (2019). Roles of ammonia-oxidizing bacteria in improving metabolism and cometabolism of trace organic chemicals in biological wastewater treatment processes: A review. Science of The Total Environment, 659, 419–441. doi:10.1016/j.scitotenv.2018.12.2

Olaniran, A. O., & Igbinosa, E. O. (2011). Chlorophenols and other related derivatives of environmental concern: Properties, distribution and microbial degradation processes. Chemosphere, 83(10), 1297–1306. doi:10.1016/j.chemosphere.2011.04

Pachapur, V. L., Sarma, S. J., Brar, S. K., Bihan, Y. L., Buelna, G., & Verma, M. (2016). Hydrogen production from biodiesel industry waste by using a co-culture of Enterobacter aerogenes and Clostridium butyricum. Biofuels, 8(6), 651–662. 10.1080/17597269.2015.1122471

Pant, S.; Kumar, A.; Ram, M. Flower pollination algorithm development: A state of art review. Int. J. Syst. Assur. Eng. Manag. 2017, 8, 1858–1866

Parham PE, Waldock J, Christophides GK, et al. Climate, environmental and socio-economic change: weighing up the balance in vector-borne disease transmission. Philos Trans R Soc B Biol Sci. 2015;370:20130551.

Peter Harvey, Sohrab Baghri, Bob Reed, Emergency Sanitation: Assessment and Program Design, Water, Engineering and Development Centre Loughborough University, Leicestershire, LE11 3TU UK, 2002. https://wedc-knowledge.lboro.ac.uk/resources/books/Emergency_Sanitation_-_Complete.pdf.

Post C, Sarala R, Gatrell C, Prescott JE (2020) Advancing theory with review articles. J Manage Stud 57(2):351– 376

Pranckute, R. Web of Science (WoS) and Scopus: The titans of bibliographic information in today’s academic world. Publications 2021, 9, 12.

Rejeb, A., Rejeb, K., Abdollahi, A., Zailani, S., Iranmanesh, M., and Ghobakhloo, M. (2021). Digitalization in food supply chains: a bibliometric review and key-route main path analysis. Sustainability 14:83. doi: 10.3390/su14010083

Reungsang, A., Sittijunda, S., & Angelidaki, I. (2013). Simultaneous production of hydrogen and ethanol from waste glycerol by Enterobacter aerogenes KKU-S1. International Journal of Hydrogen Energy, 38(4), 1813– 1825. doi:10.1016/j.ijhydene.2012.11.06

Rupasinghe, Lakmali and Pushpakumari, M. D. and Perera, G. D. N., Systematic Literature Review: On Green Innovation (December 9, 2023). Management Journal for Advanced Research Peer Reviewed and Refereed Journal, Volume-3, Issue-6, December 2023, PP. 9–21, Available at SSRN: https://ssrn.com/abstract=4660173

Sawasdee, V., Vikromvarasiri, N. & Pisutpaisal, N. Optimization of ethanol production from co-substrate of waste glycerol and acetic acid by Enterobacter aerogenes. Biomass Conv. Bioref. 13, 10505–10512 (2023). 10.1007/s13399-021-01844-9

Scarlat N, Motola V, Dallemand JF, Monforti-Ferrario F, Mofor L. Evaluation of energy potential of municipal solid waste from African urban areas. Renewable and Sustainable Energy Reviews. 2015;50(October):1269–1286. DOI: 10.1016/j.rser.2015.05.067

Tomita, A., Cuadros, D. F., Burns, J. K., Tanser, F., & Slotow, R. (2020). Exposure to waste sites and their impact on health: a panel and geospatial analysis of nationally representative data from South Africa, 2008-2015. The Lancet. Planetary health, 4(6), e223–e234. 10.1016/S2542-5196(20)30101-7

UNEP (2018). Africa Waste Management Outlook. United Nations Environment Programme, Nairobi, Kenya. www.unenvironment.org/ietc/resources (Accessed on December 11, 2024).

UNEP (United Nations Environment Programme). (2015). Global Waste Management Outlook (GWMO). UNEP DTIE International Environmental Technology Centre, Osaka. https://www.iswa.org/fileadmin/galleries/Publications/ISWA_Reports/GWMO_summary_web.pdf. (Accessed on 11 December 2024).

United Nations Environment Programme (2024). Global Waste Management Outlook 2024: Beyond an age of waste – Turning rubbish into a resource. DOI: 10.59117/20.500.11822/44939

Whiteford, R., Nurika, I., Schiller, T., & Barker, G. (2021). The white rot fungus, Phanerochaete chrysosporium, under combinatorial stress produces variable oil profiles following analysis of secondary metabolites. Journal of Applied Microbiology, 131(3), 1305–1317. doi:10.1111/jam.15013

World Health Organization (WHO), Regional Strategy for the Management of Environmental Determinants of Human Health in the African Region 2017-2021. Sixty-Seven Sessions: Victoria Falls, Republic of Zimbabwe, 28 August–1 September 2017, 2017. Provisional agenda item 9. Report of the Secretariat. WHO regional office for Africa, https://www.afro.who.int/sites/default/files/2017-12/AFR-RC67-6%20Regional%20strategy%20for%20environ%20health%20determ%20Human%20Health.pdf (Accessed on December 11, 2024).

Wu, D., Ekama, G. A., Chui, H.-K., Wang, B., Cui, Y.-X., Hao, T.-W., … Chen, G.-H. (2016). Large-scale demonstration of the sulfate reduction autotrophic denitrification nitrification integrated (SANI ®) process in saline sewage treatment. Water Research, 100, 496–507. doi:10.1016/j.watres.2016.05.052

Zeng, G. M., Chen, A. W., Chen, G. Q., Hu, X. J., Guan, S., Shang, C., Lu, L. H., & Zou, Z. J. (2012). Responses of Phanerochaete chrysosporium to toxic pollutants: physiological flux, oxidative stress, and detoxification. Environmental science & technology, 46(14), 7818–7825. 10.1021/es301006j

Zhao, J., Dong, Z., Li, J., Chen, L., Bai, Y., Jia, Y., & Shao, T. (2019). Evaluation of Lactobacillus plantarum MTD1 and waste molasses as fermentation modifier to increase silage quality and reduce ruminal greenhouse gas emissions of rice straw. The Science of the total environment, 688, 143–152. 10.1016/j.scitotenv.2019.06.236

Zhao, M.; Liu, W.; Saif, A.N.M.; Wang, B.; Rupa, R.A.; Islam, K.M.A.; Rahman, S.M.M.; Hafiz, N.; Mostafa, R.; Rahman, M.A. Blockchain in Online Learning: A Systematic Review and Bibliographic Visualization. Sustainability 2023, 15, 1470. 10.3390/su15021470

Zhong, Q., Wang, L., and Cui, S. (2021). Urban food systems: a bibliometric review from 1991 to 2020. Foods 10:662. doi: 10.3390/foods10030662

Li, M., Zhan, A., Rahman, T. T., Jiang, T., & Hou, L. (2025). From wastewater to resistance: characterization of multidrug-resistant bacteria and assessment of natural antimicrobial compounds. Frontiers in Microbiology, 16. 10.3389/fmicb.2025.1612534

Yang, M., Chen, B., Xin, X., Song, X., Liu, J., Dong, G., Lee, K., & Zhang, B. (2020). Interactions between microplastics and oil dispersion in the marine environment. Journal of Hazardous Materials, 403, 123944. 10.1016/j.jhazmat.2020.123944

